# Bark-associated diazotroph communities are a cryptic source of nitrogen in forests

**DOI:** 10.1101/2025.11.14.688567

**Authors:** Paula Gomez-Alvarez, Chuwen Zhang, Luke C. Jeffrey, Pok Man Leung, Perran L.M. Cook, Dirk Erler, Anthony J. Kohtz, Johannes Dittmann, Wei Wen Wong, Scott Cramb, Sebastian Euler, Montgomery Hall, Billie Hopkins, Chris Lee, Scott G. Johnston, Xiyang Dong, Chris Greening, Damien T. Maher

**Affiliations:** Faculty of Science and Engineering, Southern Cross University, East Lismore, NSW 2480, Australia; Key Laboratory of Marine Genetic Resources, Third Institute of Oceanography, Ministry of Natural Resources, Xiamen 361005, China; Department of Microbiology, Biomedicine Discovery Institute, Monash University, Clayton, VIC 3800, Australia; Water Studies, School of Chemistry, Monash University, Clayton, VIC 3800, Australia; School of Biological Sciences, Monash University, Clayton, VIC 3800, Australia

**Author notes:** Corresponding authors: Prof Chris Greening, Prof Damien Maher, Paula Gomez-Alvarez. These authors substantially and equally contributed to this study (Chuwen Zhang, Luke C. Jeffrey, Pok Man Leung).

## Abstract

Nitrogen is an essential nutrient limiting forest productivity. Plants cannot access atmospheric N_2_ directly and rely on diazotrophic bacteria to fix nitrogen into bioavailable forms such as ammonium. Within forests, biological nitrogen fixation (BNF) occurs primarily in soils and root nodules. However it is unclear whether the extensive microbial communities recently discovered in tree bark can also fix nitrogen. Here we combine field measurements, metagenomic profiling, and biogeochemical assays to show that bark-dwelling diazotroph communities are abundant and active across diverse tree species. Bark from eight Australian tree species showed exceptionally high C:N ratios and depleted nitrogen stable isotope signatures (δ^15^N) consistent with locally fixed nitrogen, suggesting strong selection for diazotrophs. Consistently, bark microbial communities harbour phylogenetically and physiologically diverse diazotrophs, averaging ∼10^12^ cells m^-2^ and in higher relative abundance than underlying soils. Canonical and alternative nitrogenases were detected across eight bacterial phyla, primarily bark-adapted Alphaproteobacteria, Acidobacteriota, and Verrucomicrobiota, with strong signatures of purifying selection. Stable isotope labelling experiments demonstrated that bark-dwelling diazotrophs fix nitrogen at rates varying with tree species and habitat. In line with the presence of methane-oxidising diazotrophs, BNF was strongly stimulated by methane addition and suppressed by methanotroph inhibitors. Initial upscaling suggests bark-associated nitrogen fixation contributes up to 3.8 Tg N yr^-1^ globally (∼6% of natural terrestrial BNF), with further studies required to constrain this budget and its contribution to tree nitrogen demands. Altogether, this discovery of substantial above-ground nitrogen inputs revises our understanding of forest nutrient cycles and redefines the functional scope of the caulosphere.

## Introduction

Despite being the most abundant element in the atmosphere, nitrogen (N) limits forest productivity and carbon sequestration^1,2^. Plants depend on the conversion of atmospheric nitrogen into bioavailable ammonia^3^ by diazotrophic microorganisms through the process of biological nitrogen fixation (BNF)^4–6^. Catalysed by the nitrogenase enzyme complex^7^, BNF requires a substantial energy and reductant supply, with at least 16 ATP consumed to fix each N_2_ molecule^8^. Within forests, BNF has traditionally been attributed root-associated symbionts, with free-living diazotrophs in soil, litter, and increasingly the phyllosphere have been also recognised^5,9–12^. For example, a range of diazotrophs inhabit tree leaves, including methylotrophic Alphaproteobacteria and photosynthetic Cyanobacteria, though they typically fix nitrogen at low rates (5.0 ± 1.2 ng N g^-1^ day^-1^) and hence only fractionally contribute to nitrogen input^13^. Comprehensive characterisation of forest diazotrophic communities is critical for consolidated nitrogen budgets and modelling forest productivity amid environmental change, especially since BNF may limit forest responses to rising carbon dioxide levels^14–17^.

Tree stems are extensive and persistent above-ground habitats, yet remain largely unexplored as sites for BNF. These tissues accumulate plant-derived polymers and transport water, gases, and nutrients, creating environments conducive for microbial growth and biogeochemical activities^18–20^. For example, recent studies have revealed that bark (i.e. the caulosphere) harbours abundant, unique, and active microbial communities with globally significant roles in climate-active gas cycling^20–23^. Nitrogen fixation genes have been detected in the bark of a few tree species, including poplar, willow, and avocado, sometimes at levels comparable to or exceeding those in surrounding soils^24–26^. There are also historical reports of nitrogen fixation, based on the acetylene reduction assay, in stem tissues^27–30^. Yet the identity, capabilities, and direct activities of these bark-dwelling diazotrophs remain to be demonstrated. Given there are over three trillion trees worldwide^31,32^ and bark spans a surface area comparable to the global land surface area^22^, bark-associated diazotrophs may constitute a major previously unrecognised source of bioavailable nitrogen. Here, we combined genome-resolved metagenomics with ^15^N-N₂ isotope tracing to characterise bark-dwelling diazotroph communities and demonstrate they are prevalent and actively fix nitrogen across eight tree species from four forest types. These results necessitate a reevaluation of the current paradigm around N cycling in forest ecosystems.

## Results

### High C:N ratios and nitrogen stable isotope signatures suggest BNF is active in tree bark

We sampled bark, leaves, leaf litter and underlying soil from three individual trees of eight tree species across Australian wetland, upland, heathland, and mangrove forests in northern New South Wales, Australia **(Fig. S1; Table S1)**. As expected, bark carbon content was significantly higher than the underlying soil and leaf litter (49.9 ± 0.9% bark; 9.5 ± 1.9% soil; Dunn’s test, *p* < 0.0001), while nitrogen content was comparable between bark and soil (0.39 ± 0.05% and 0.40 ± 0.07%, respectively; *p* = 1.00) **(Table S2, S3)**. This resulted in exceptionally high bark C:N ratios (169.9 ± 16.9), compared to soil (22.9 ± 1.2), leaf litter (57.0 ± 5.9), and leaves (25.4 ± 2.7) (all *p* < 0.005, **Fig. 1a, Table S2 – S5**), suggesting that microbes residing within tree bark are likely strongly nitrogen-limited and may rely on N_2_ fixation to meet anabolic demands. To evaluate this hypothesis, we measured the nitrogen stable isotope signatures (δ^15^N) in bark samples. Depleted δ^15^N values in nitrogen pools typically indicate nitrogen fixation as the source, since diazotrophs assimilate atmospheric N_2_ with a δ^15^N typically between -2 and +1‰^33,34^. Seven of the tree species exhibited depleted bark δ^15^N values (-1.20 ± 0.22‰), consistent with nitrogen derived primarily from BNF^34^ **(Fig. 1b, Table S2)**. The mangrove species *Avicennia marina* was a notable exception, exhibiting significantly elevated δ^15^N (+4.48 ± 0.02‰; Tukey’s HSD, *p* < 0.0001), reflecting assimilation of remineralised inorganic nitrogen (e.g. NO_3_^-^) from marine environments^35,36^, over atmospheric nitrogen fixation **(Fig. 1b)**. Importantly, the bark δ^15^N values were lower than soil δ^15^N (**Tables S2-3**, *p* < 0.001), suggesting that nitrogen in tree bark from terrestrial species is likely derived from recent endogenous nitrogen fixation. However, the relative contributions from root- or bark-associated diazotrophs to these signatures remain unresolved.

**Figure 1.**
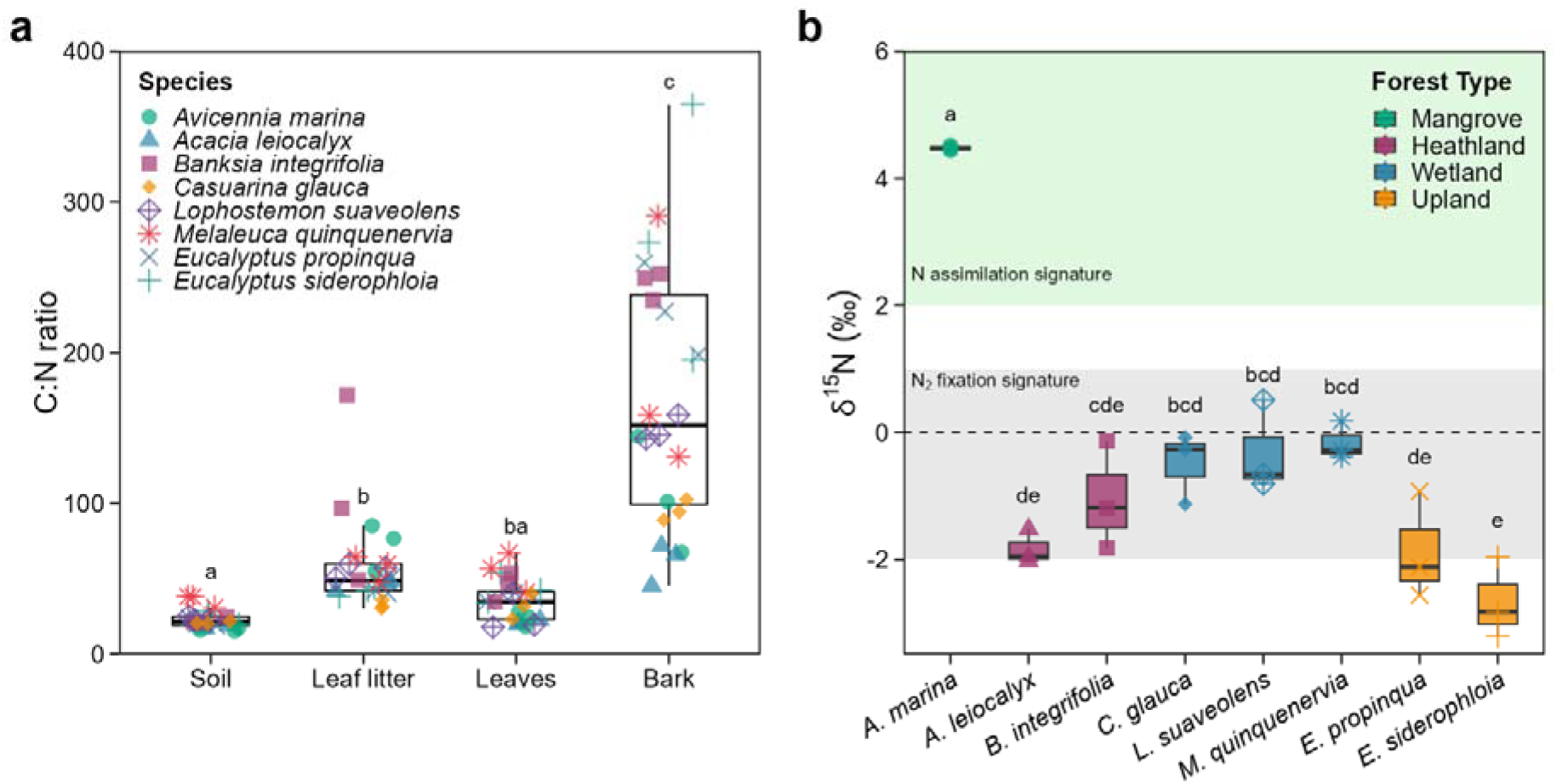
Nitrogen elemental ratios and isotopic signatures in tree bark suggest biological nitrogen fixation. **(a)** Carbon-to-nitrogen ratios (C/N) for four forest compartments (bark, leaves, leaf litter and soil) associated with eight tree species (n = 3 trees). Different symbols and colours are used for each tree species. Boxplots show median (central line), interquartile range (box), and 1.5× interquartile range (whiskers). C/N ratios were significantly different between forest compartments (Kruskal-Wallis χ^2^ = 71.3, df = 3, *p* < 0.0001), with bark exhibiting exceptionally high ratios compared to soil (p < 0.00001), leaf litter (p = 0.0085) and leaves (p < 0.00001) indicating severe nitrogen limitation in the bark. Letters denote significant pairwise differences (Dunn’s test with Bonferroni adjustment) **(b)** Nitrogen stable isotope signatures (δ^15^N (‰)) in bark samples across eight tree species (n = 3 trees) grouped by forest type. Shaded regions indicate isotopic signatures typically associated with nitrogen assimilation (green)^36^ and nitrogen fixation (grey)^33,34^. Letters indicate significant differences between species (one-way ANOVA/Tukey’s HSD).

### Phylogenetically and physiologically diverse diazotrophs are abundant in tree bark

We next sequenced bark metagenomes from the 24 sampled trees to investigate whether their microbial communities could mediate BNF **(Table S6)**. Consistent with recent studies^20,37^, the bark of every tree hosted abundant, diverse, and functionally specialised microbial communities with high capacities for both aerobic and anaerobic metabolism, carbohydrate degradation, and trace gas cycling **(Table S6)**. In line with nitrogen limitation, nitrification genes were generally absent and denitrification genes were 3 – 5 fold lower in bark than surrounding soil communities. By contrast, based on the nitrogenase marker gene (*nifH*), bark-associated diazotrophs were abundant in the seven terrestrial tree species, comprising on average 21.5 ± 12% of the microbial community; their abundance varied significantly across tree species, from 43.4% in *Casuarina glauca* to 13.3% in *Banksia integrifolia* **(Fig. 2a)**. The relative abundance of diazotrophs within terrestrial trees was significantly higher in tree bark than adjacent soils (8.5 ± 5.5%; *p* = 0.011) **(Fig. 2a)** and moderately correlated with bark δ^15^N (*R^2^* = 0.19; **Fig. S2**). The mangrove species, by contrast, showed the opposite trend, though diazotrophs remain abundant in bark (17.6 ± 6.1%) **(Table S6)**. Based on qPCR quantification of the 16S rRNA gene copy numbers, we estimate diazotrophs vary in abundance from 1.7 billion and 12.5 trillion diazotrophic bacteria per square metre of tree bark across the eight species **(Table S6)**. It should be noted that, although diazotrophs were especially enriched in tree bark, surrounding soils still showed a high capacity for diazotrophy and are likely a major source of the fixed nitrogen for each tree.

**Figure 2.**
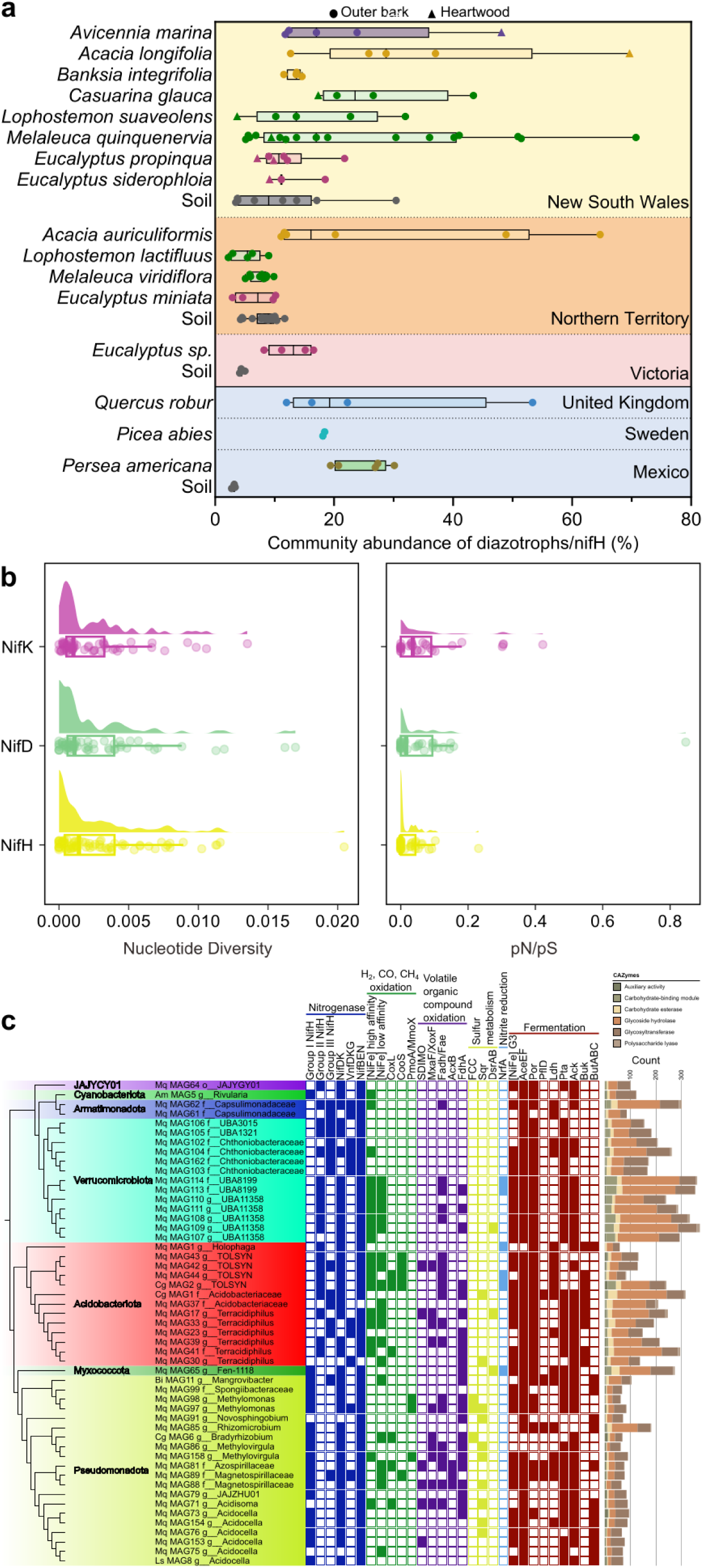
Nitrogenase genes are widespread and under purifying selection in tree bark microbial communities. **(a)** Boxplot showing abundance of nitrogenase genes (*nifH*) within microbial communities of different tree species and between outer bark, heartwood, and soil across different locations. **(b)** Nucleotide diversity and ratio of nonsynonymous to synonymous substitutions (pN/pS) in the genes (*nifH*, *nifD*, *nifK*) encoding the three nitrogenase structural subunits. **(c)** Phylogenetic diversity and metabolic capabilities of the 51 nitrogenase-encoding MAGs. Genes present in each MAG are shaded. Abbreviations: NifH (nitrogenase reductase), NifDK ([MoFe]-nitrogenase catalytic subunits), VnfDKG (vanadium nitrogenase catalytic subunits), NifBEN (nitrogenase iron-molybdenum cofactor biosynthesis proteins), [NiFe] high affinity (high-affinity uptake [NiFe]-hydrogenase), [NiFe] low affinity (low-affinity uptake [NiFe]-hydrogenase), CoxL (form I [MoCu]-CO dehydrogenase), CooS (anaerobic [NiFe]-CO dehydrogenase), PmoA (particulate CH_4_ monooxygenase), MmoX (soluble CH_4_ monooxygenase), SDIMO (soluble diiron monooxygenases), MxaF (calcium-dependent methanol dehydrogenase), XoxF (lanthanide-dependent methanol dehydrogenase), Fadh (formaldehyde dehydrogenase), Fae (formaldehyde-activating enzyme), AcxB (acetone carboxylase), FdhA (formate dehydrogenase), FCC (flavocytochrome *c* sulfide dehydrogenase), Sqr (sulfide:quinone oxidoreductase), DsrAB (dissimilatory sulfite reductase), NrfA (ammonifying nitrite reductase), [NiFe] G3 (group 3 [NiFe]-hydrogenase), AceEF (pyruvate dehydrogenase complex), Por (pyruvate:ferredoxin oxidoreductase), PflD (pyruvate-formate lyase), Ldh (lactate dehydrogenase), Pta (phosphate acetyltransferase), Ack (acetate kinase), Buk (butyrate kinase), and ButABC (butanediol dehydrogenase).

To better understand the nature and capabilities of the bark-associated diazotrophs, we analysed 209 species-level metagenome-assembled genomes (MAGs) reconstructed from the eight Australian tree species **(Table S4)**. Fifty-one MAGs (i.e. 24%, in line with the short-read data) encoded nitrogenase structural and/or biosynthesis operons. These spanned eight different phyla and dominant bark-associated genera, such as *Terracidiphilus*, *Acidocella*, Pedosphaerales UBA11358, and *Methylomonas*^20,21,37^, as well as a cyanobacterium (*Rivularia*), a rhizobium (*Bradyrhizobium*), and the candidate phylum JAJYCY01 **(Fig. 2d)**. Read mapping revealed that several diazotrophic MAGs (e.g. *Terracidiphilus*, *Acidocella, Mangrovibacter*, and *Bradyrhizobium*) were highly abundant in different tree species, for example with one *Acidocella* MAG comprising 26.5% of the bark community of a *Melaleuca* tree **(Table S4)**. This suggests nitrogen fixation may be a key adaptive strategy for bark inhabitants to grow amid nitrogen limitation. Also consistent with strong purifying selection pressure, analyses of evolutionary indices in the nitrogenase structural genes (*nifHDK*) revealed low nucleotide diversity **(Fig. 2b)** and exceptionally low nonsynonymous to synonymous polymorphism ratios (mean pN/pS = 0.006 *nifH*, 0.022 *nifD*, 0.039 *nifK*; **Fig. 2b)**. This pronounced signal of purifying selection reflects the structural and mechanistic complexity of nitrogenase, especially the dinitrogenase reductase NifH, which is highly conserved and intolerant to amino acid substitutions that disrupt electron transfer, ATP hydrolysis, or interactions with its dinitrogenase protein partners^8,38^.These data indicate that diazotrophs depend on a functional nitrogenase complex to adapt to nutrient-limited bark environments.

Phylogenetic analysis of NifH protein sequences retrieved from 46 MAGs **(Fig. 3)** and 242 unbinned contigs **(Fig. S3)** revealed substantial structural and functional diversity. Most bark nitrogenases were affiliated with canonical molybdenum-containing nitrogenases. Group I (44% sequences) nitrogenases were encoded primarily by Alphaproteobacteria, as well as some Myxococcota and Actinomycetota MAGs, while group II (34% sequences) were encoded by members of the Acidobacteriota, Verrucomicrobiota, and candidate phylum JAJYCY01. Alternative vanadium/iron nitrogenases (group III; 19% sequences) were also distributed across multiple phyla, with several unbinned canonical group IVa nitrogenases also detected **(Fig. 3, Fig. S3)**. Several MAGs, including three species each of Acidobacteriota, Verrucomicrobiota, and Alphaproteobacteria, co-encoded both molybdenum and vanadium/iron nitrogenases, suggesting metabolic flexibility to continue fixation amid fluctuating micronutrient availability. Gene neighbourhood analyses showed that *nifH* clustered within operons containing the nitrogenase structural genes (*nifD*, *nifK*) in diverse orientations, typically accompanied by nitrogenase biosynthesis (*nifENB*), regulatory (e.g. *nifL*_1_ and *nifL*_2_), and accessory genes **(Fig. 3)** Nitrogenases reconstructed from surrounding soil and water metagenomes formed distinct phylogenetic clades to those from the bark metagenomes, consistent with most bark-associated diazotrophs being resident, rather than environmentally-transferred transient, community members.

**Figure 3.**
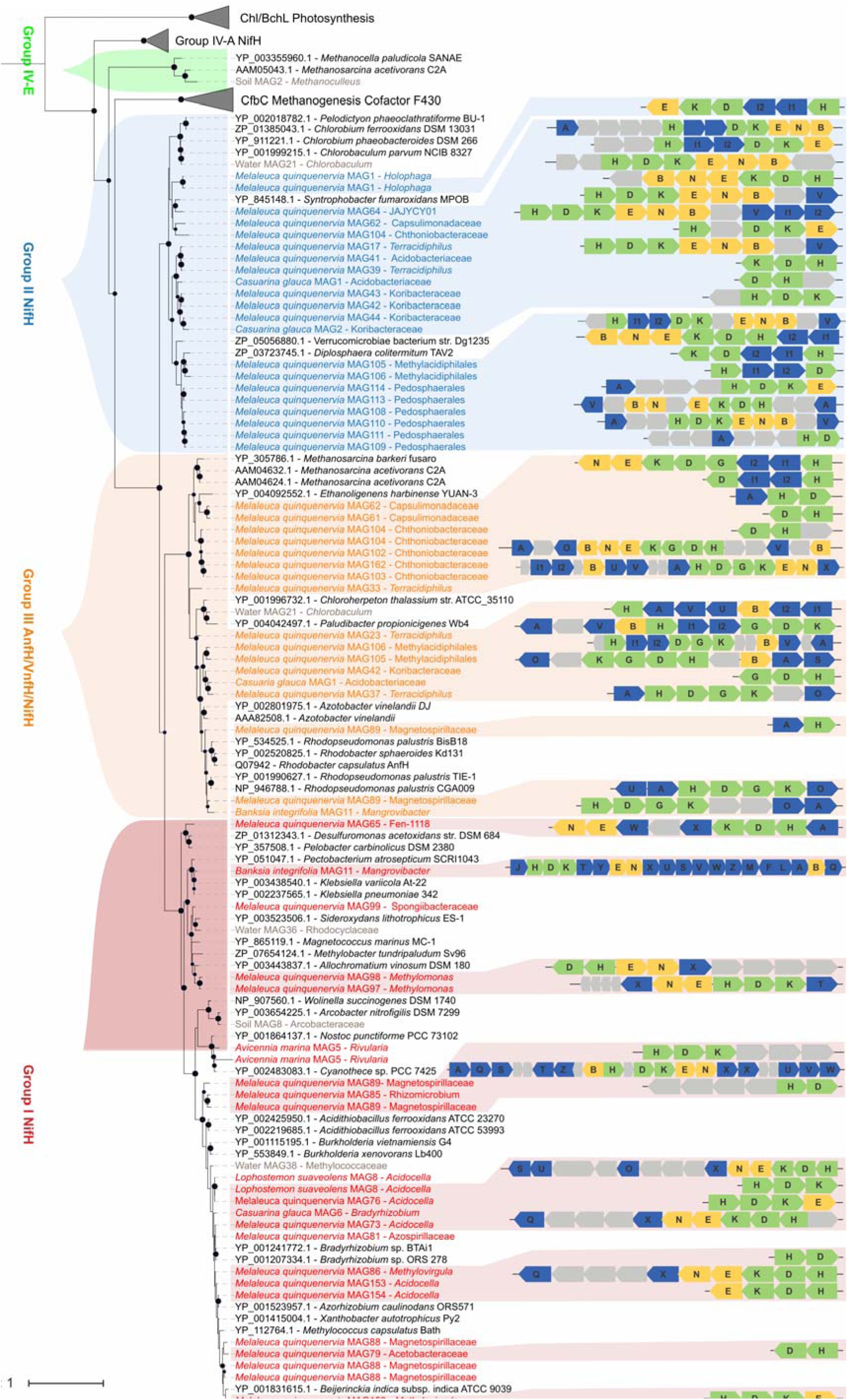
Phylogenetic tree and genetic organisation of the diverse nitrogenases encoded by tree bark microbial communities. The tree was constructed based on NifH/VnfH/AnfH protein sequences. The scale bar represents one amino acid substitution per sequence position. The bootstrap values of >70% are indicated as black circles at the nodes, and scale bars indicate the mean number of substitutions per site. Branch colours indicate distinct clades of bark-associated nitrogenases, while branches corresponding to nitrogenases originating from surrounding soil and aquatic metagenomes are shown in grey.

We further analysed the metabolic capabilities of the 51 bark-associated diazotrophs based on their relatively complete genomes (average 86% complete) **(Table S4, Fig. 2c)**. In agreement with our recent analyses^20^, most taxa including the dominant Alphaproteobacteria, Verrucomicrobiota, and Acidobacteriota are predicted to be facultative anaerobes, capable of aerobic respiration (100%), acetate fermentation (59%), and fermentative hydrogen production (75%). Although most are predicted to mediate organoheterotrophic growth using plant-derived carbohydrates and volatile organic compounds, six MAGs are predicted to fix CO_2_ using RuBisCO, all chemosynthetically except the cyanobacterium. Bark-associated diazotrophs showed widespread capacity to gain energy by consuming H_2_ (55% MAGs), formate (65% MAGs), methanol (21% MAGs), CO (15% MAGs), and sulfide (24% MAGs), including high-affinity enzymes to scavenge atmospheric trace gases. Importantly, two MAGs from the *Methylomonas* and *Methylovirgula* were predicted to be capable of methanotrophy; whereas the former is a widespread bark lineage shown to fix nitrogen in culture-based studies^39^, the latter was originally reported as a facultative methylotroph but includes several methanotrophic isolates^40^. Together, these findings suggest bark-associated diazotrophs can couple the energy demanding process of nitrogen fixation to the use of diverse energy sources, including organic plant compounds, high-energy gases and volatiles, and occasionally light, under a range of environmental conditions.

### Bark-associated diazotrophs mediate nitrogen fixation and are stimulated by methane

To directly test whether bark-associated microbial communities actively fix nitrogen, we quantified BNF rates across eight tree species using ^15^N-N_2_ isotope tracing assays. The use of ^15^N-N_2_ tracing provides direct high-resolution measurements of nitrogen incorporation, is robust against potential inhibition of microbial activity associated with acetylene reduction assay (ARA)^41^, preserves the natural structure of bark, and accounts for inter-individual variation in baseline δ^15^N among trees. To account for within- and between-tree variability, we sampled three trees for each species and incubated triplicate bark samples of each tree **(Table S5)**. Nitrogen fixation was detected in all tree species, providing the first direct evidence that bark-associated microbes contribute newly fixed nitrogen to trees **(Fig. 4a)**.

**Figure 4.**
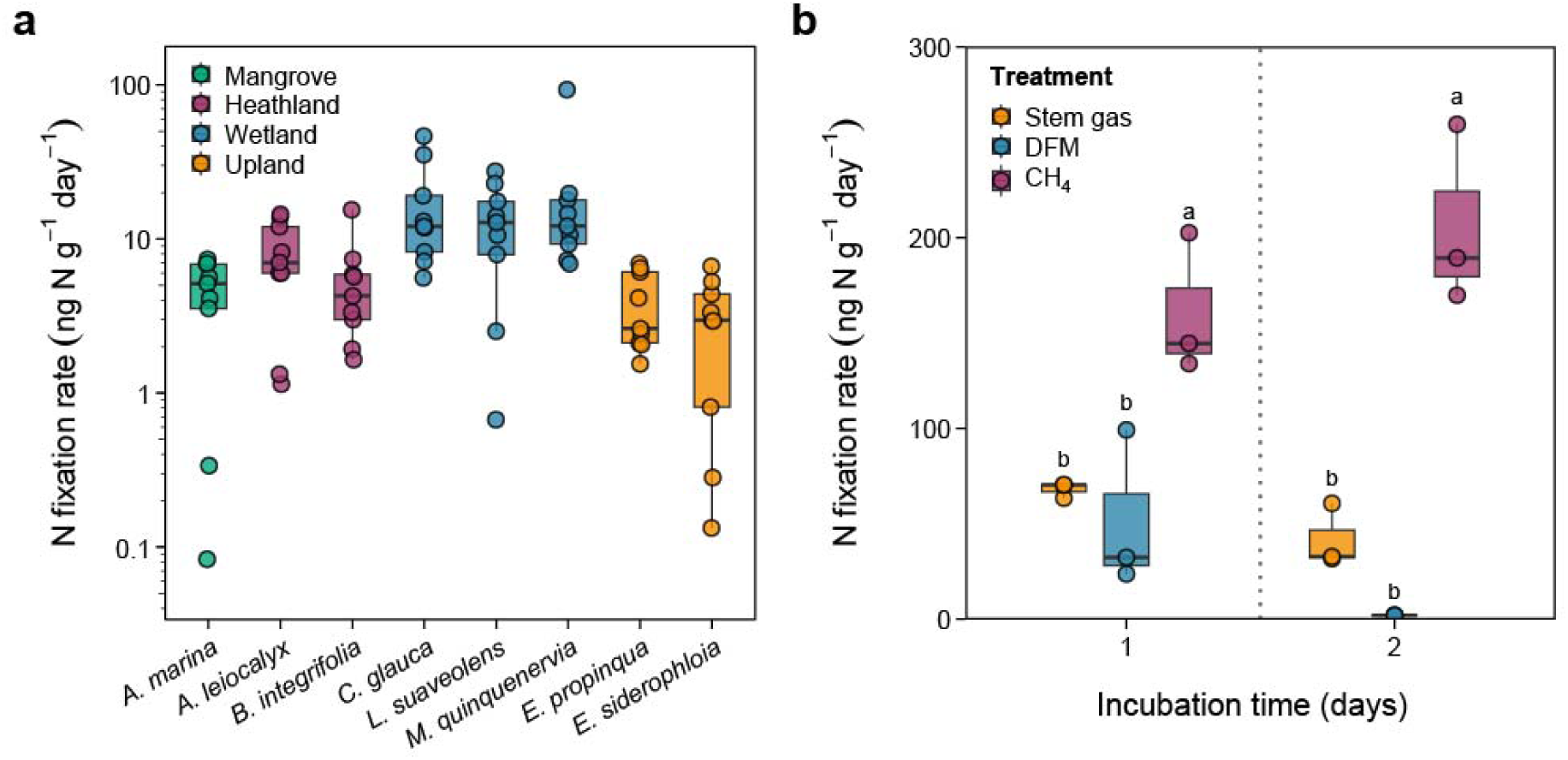
Biological nitrogen fixation occurs across diverse tree species. **(a)** Nitrogen fixation rates measured with the ^15^N_2_ assimilation method in triplicate bark samples from three individual trees of 8 different species grouped by habitat (n = 9). The y-axis is shown on a log_10_ scale to enhance visualisation of differences between species. Nitrogen fixation rates differed significantly among species (LME with box-cox transformation λ = 0.222; main effect of species χ^2^ = 51.4, df = 8, *p* < 0.001), while variability between trees of the same species was negligible (SD = 5.66 × 10^-5^). BNF rates for coastal wetland species exceeded upland eucalyptus and *Avicennia marina* (*p* < 0.05), *M. quinquenervia* exceeded *Banksia integrifolia* (*p* = 0.009) and *Acacia leiocalyx* (*p* = 0.047), and *A. leiocalyx* exceeded *E. siderophloia* (*p* = 0.047). **(b)** Effect of adding the methanotrophy substrate methane (0.26% CH_4_) and the methanotrophy inhibitor difluoromethane (DFM) on bark-associated BNF in *M. quinquenervia* measured with ^15^N_2_ incorporation assays. Bark samples were incubated in triplicate with (field-collected) stem gas (naturally containing 655 ppm CH_4_), stem gas with 0.26% CH_4_ (2,600 ppm; final CH_4_ concentration 3,060 ppm) stem gas + 2% DFM over one and two days. Treatment had a significant effect on nitrogen fixation (two-way ANOVA: F_2,12_ = 44.9, *p* < 0.0001), with a significant interaction with incubation time (F_2,12_ = 4.01, *p* = 0.046). Methane addition significantly increased nitrogen fixation rates compared to the use of stem gas at both Day 1 (Tukey’s HSD, *p* = 0.008) and Day 2 (*p* = 0.0001), while DFM-treated samples exhibited significantly lower rates than methane supplemented samples (Day1: *p* < 0.0025; Day 2: *p* < 0.0001). Letters denote significant differences between treatments (two-way ANOVA/Tukey’s HSD).

BNF rates strongly varied with tree species and environment **(Fig. 4a; Fig. S4)**. Consistent with expectations from δ^15^N signatures and metagenomic profiling, freshwater wetland trees exhibited the highest BNF activity. *M. quinquenervia* averaged 21.3 ± 9.1 ng N g^-1^ day^-1^, while *Lophostemon suaveolens* (12.9 ± 2.9 ng N g^-1^ day^-1^) and *C. glauca* (17.6 ± 4.7 ng N g^-1^ day^-1^) also supported high nitrogen fixation. Both heathland species *A. leiocalyx and B. integrifolia* exhibited moderate nitrogen fixation (averaging 7.8 and 5.4 ng N g^-1^ day^-1^ respectively), whereas lower fixation was observed for the two upland eucalypts and *A. marina*, the latter in line with the δ^15^N inferences that this mangrove species likely assimilates remineralised nitrogen rather than recently fixed nitrogen **(Fig. 4a)**. Linear mixed-effects modelling confirmed that BNF rates differed significantly among species and forest types (χ^2^_8_ = 51.4, *p* < 0.001; AIC = 268.35, BIC = 292.43), with model diagnostics confirming assumptions were met, and cross-species differences were stronger than tree-to-tree variation within species (random-effect variance = 3.2 × 10^-9^, SD = 5.7 × 10^-5^). Yet substantial variation in BNF rates were observed both between trees of the same species and among bark subsamples within individual trees **(Fig. S4)**. This was particularly pronounced for *M. quinquenervia*, where some samples showed relatively low activity and others reached the maximum observed rate of 93.1 ng N g^-1^ day^-1^ **(Table S8)**. The variability between replicates of the same species may reflect variability in physicochemical properties and microbial communities in different bark sections, including potentially the importance of microsites where optimal oxygen levels, temperature, moisture, and energy availability converge. Altogether, these patterns suggest that bark nitrogen fixation is widespread but highly species- and microhabitat-dependent.

Given *M. quinquenervia* bark communities exhibited the highest and most variable BNF rates, we explored whether fluctuations in methane availability could contribute to this variability. Our previous studies show that methanotroph abundance and activity varies by an order of magnitude between *M. quinquenervia* wetland trees, comprising up to a quarter of the bacterial community^20,21^. In laboratory incubations, BNF occurred in the presence of field-collected stem gas (containing 655 ppm CH_4_) and rates were stimulated by 2.4-fold (*p* = 0.008) and 4.9-fold (*p* = 0.0001) after one and two days of stimulation with 2600 ppm methane respectively (**Fig. 4b, Table S9**). In contrast, administration of the specific methane monooxygenase inhibitor difluoromethane^42^ (2% DFM) strongly reduced activity (*p* < 0.0001). These results are consistent with metagenomic predictions that methanotrophic diazotrophs are abundant in *M. quinquenervia* bark and may use ATP generated from methane oxidation to drive nitrogen fixation. Collectively, these findings validate that methanotrophs provide a direct and dynamic source of fixed nitrogen in bark communities and that nitrogen fixation activity is strongly and dynamically regulated by methane availability.

### Bark-associated diazotrophs are globally distributed and may contribute to nitrogen budgets

To evaluate whether these findings are translatable across a wider spatiotemporal scale, we further sampled and sequenced metagenomes of bark and heartwood of the same eight tree species in two seasons of a different year. The results suggested that tree-associated diazotrophs are stable across seasons, and are abundant in both and bark **(Fig. 2a, Fig. S5a)**. Moreover, diazotrophs were similarly abundant in metagenomes of five tree species from tropical forests (Northern Territory, AU) and temperate upland forests (Victoria, AU) (**Fig. 2a**), across the full height of a tree (**Fig. S5b**) as well as published metagenomes of three species sampled outside Australia, namely English oak (*Quercus robur*)^43^, Hass avocado (*Persea americana*)^44^, and Norway spruce (*Picea abies*)^19^ (**Fig. 2a**). This indicates that bark associated BNF is a globally significant process across diverse forest types. Finally, we performed an upscaling analysis based on the average measured BNF (0.0189 g N m^-2^ yr^-1^) and the estimated global woody surface area (84 – 206 million km^2^)^22^. This yields an annual flux of approximately 1.5 – 3.8 Tg N yr^-1^ from bark-associated BNF, corresponding to 2.3% to 5.8% of the estimated natural terrestrial BNF (65 Tg N yr^-1^)^45^. Combined, our field measurements, biogeochemical assays, and metagenomic profiling provide multiple lines of evidence that bark diazotrophs globally and significantly contribute to nitrogen fixation, though much wider sampling is required to constrain their contribution to global nitrogen budgets.

## Discussion

This discovery of abundant, diverse, and active diazotrophs in tree bark reframes our understanding of plant-microbe interactions and forest nutrient inputs. Across distinct forest and tree types, the extreme carbon richness and nitrogen scarcity in bark impose strong selective pressure on microbial communities, opening critical ecological roles to be filled by resident nitrogen-fixing bacteria. The abundance and diversity of diazotrophs across multiple phyla, each maintaining highly conserved nitrogenase genes under strong purifying selection, indicates that nitrogen fixation is not incidental but rather a fundamental adaptive trait required for microbial communities inhabiting the bark environment. In this regard, bark appears to harbour self-sustaining microbial ecosystems that can continually meet nitrogen needs, while using both atmospheric trace gases and volatile organic compounds as energy and carbon supplies^20^. Yet these bark-associated communities remain connected to wider biogeochemical processes through their interactions with the host tree and the forest floor.

Energetically, the high ATP demands of bark-associated diazotrophy appear to be met by a mosaic of locally available energy sources. Metagenomic evidence indicates that diazotrophs exploit not only plant-derived carbohydrates, but also variable volatile organic compounds (e.g. methanol, formaldehyde), atmospheric trace gases (CH_4_, H_2_, CO), sulfide, and light, to meet energy needs under both oxic and anoxic conditions. Such metabolic flexibility suggests bark communities can simultaneously or alternately use various pathways to meet the substantial energetic demands of nitrogen fixation under fluctuating environmental conditions. Indeed, with atmospheric hydrogen oxidation yielding up to 2 ATP molecules per H_2_ molecule oxidised^46^, even trace gases may provide sufficient energy for nitrogenase activity. This diversity of energy coupling links bark BNF to broader forest carbon, hydrogen, and sulfur cycling. Particularly significant is the observation that nitrogen fixation is coupled to methane oxidation. Given wetland trees filter methane emissions and upland trees are a major net sink of atmospheric methane^20–22^, this coupling represents a previously unrecognised feedback between the nitrogen and carbon cycles. Methanotrophic diazotrophy may therefore serve as both a methane sink and a local nitrogen source, reinforcing the self-sufficiency of bark associated microbial communities while influencing greenhouse gas exchange.

The contribution of bark-associated free-living diazotrophs to global nitrogen budgets remains poorly constrained. Our measurements suggest an average nitrogen fixation rate of 0.0189 g N m^-2^ yr^-1^, scaling to 1.5 – 3.8 Tg N yr^-1^ globally, suggesting bark-associated BNF makes a significant and overlooked contribution to natural terrestrial biological nitrogen fixation^45^. Bark-associated BNF may be disproportionately important in N-limited ecosystems where it provides direct and localized inputs across plant surfaces. The large uncertainties inherent in this global extrapolation – stemming from limited geographic representation, climatic variation, and species-specific differences – underscore the need for expanded measurements across biomes to better constrain this previously overlooked component of the terrestrial nitrogen cycle. Nitrogen fixed within bark may enter the host tree through exchange with xylem or phloem tissues, in line with the presence of diazotrophs in heartwood. It may also be released to soils via bark stem flow^47^, or as bark exfoliates, a particularly common process in *Melaleuca* and *Eucalyptus* species. These pathways suggest that bark-associated diazotrophy, while locally self-sustaining, could contribute nitrogen both to the tree and to surrounding ecosystems. However, the magnitude and frequency of such transfers remain unknown, and soil- and root-associated symbiotic diazotrophs likely remain the main contributor to tree nitrogen supply. Altogether, bark-associated BNF represents a substantial and cryptic component of the global nitrogen cycle. Future research quantifying nitrogen transfer pathways, temporal dynamics across seasons and variability across global forest biomes will be essential to integrate bark-associated BNF into ecosystem and Earth system models.

## Materials and Methods

### Study sites and forest types

Bark, heartwood, soil, leaf litter and leaf samples were collected from trees spanning four different Australian forest ecosystems in northern New South Wales: (i) coastal freshwater wetlands dominated by *Melaleuca quinquenervia* (broad-leaved paperbark), *Lophostemon suaveolens* (Swamp Box) and *Casuarina glauca* (Swamp Oak); (ii) coastal mangrove forests inhabited by *Avicennia marina* (Grey Mangrove); (iii) subtropical upland forests with *Eucalyptus siderophloia* (Iron bark) and *Eucalyptus propinqua* (Small-fruited Grey Gum); and (iv) coastal heathlands containing *Banksia integrifolia* (Coastal Banksia) and *Acacia leiocalyx* (Black Wattle). To expand the geographic scope and the survey of bark-associated diazotrophs through metagenomic analysis, additional bark and soil samples were collected from Australian tropical wetland forests (Territory Wildlife Park, Darwin, Northern Territory) and temperate upland forests (Kurth Kiln Regional Park, Victoria) **(Figure S1)**.

### Sample collection and sequencing campaigns

Bark, heartwood and soil samples were collected using sterile techniques to minimise contamination. Approximately 10 × 12 cm bark swatches were collected from three trees of each species at a consistent lower stem height (mean 83.7 ± 4.2 cm above soil or wetland water level) using a sterile chisel and a mallet. Heartwood samples were collected using a sterile increment borer (5.15 mm diameter) inserted to a depth of 3 – 5 cm into the sapwood. The borer was sterilised with 70% ethanol and flame-sterilised between samples. Paired soil samples (0-5 cm depth) were collected below each tree. Samples were stored on ice during transport, then at -80°C until DNA extraction. For the subtropical northern New South Wales sites, preliminary bark (n = 7 trees) and soil (n = 2) sampling from *M. quinquenervia* was conducted for metagenomic sequencing on May 20, 2020, followed by a wet and a dry campaign that included bark and heartwood of eight tree species on May 10, 2021 and October 19, 2021, respectively, and a final comprehensive sampling campaign of bark and the surrounding soils from all eight tree species (n = 3 trees per species) on August 10, 2022 **(Figure S1, Table S1)**. Sampling at temperate Kurth Kiln Regional Park was conducted on March 22, 2023, in which four bark samples were collected from two dominant eucalypts, peppermint (*Eucalyptus dives* / *Eucalyptus radiata*) and stringybark (*Eucalyptus baxteri* / *Eucalyptus obliqua*), in addition to three soil samples. The tropical sampling campaign at the Territory Wildlife Park (Darwin, Northern Territory) was conducted on April 2-3, 2025, where bark samples were collected at 0.6 m height from four tree of each of three representative tropical wetland species (*Melaleuca viridiflora, Acacia auriculiformis* and *Lophostemon lactifluus*) and one upland eucalypt species (*Eucalyptus miniata*) (**Figure S1**).

### DNA extraction and sequencing

For the subtropical NSW and tropical NT samples, 0.1 – 0.2 g samples of bark were snap frozen in liquid nitrogen and ground into fine powder. Genomic DNA was extracted using the Synergy 2.0 Plant DNA extraction kit (OPS Diagnostics) and the DNeasy PowerSoil kit (Qiagen), with negative controls (blank extractions using PCR-grade water) included in each batch to monitor for contamination. Soil DNA was extracted from 0.25 g of samples using the DNeasy PowerSoil kit (Qiagen). DNA quantity and quality were assessed using a Qubit fluorometer and NanoDrop ND-1000 spectrophotometer. Samples with quantifiable DNA were sent to the Australian Centre for Ecogenomics, University of Queensland for metagenomic sequencing. Metagenomic shotgun libraries were prepared using the Nextera XT DNA Library Preparation Kit (Illumina) and passed to paired-end sequencing (2 × 150 bp) on a NovaSeq 6000 platform (Illumina). For the temperate bark samples, 0.03 – 0.2 g of material was ground prior to extraction using a mixer mill (Retsch MM400, Retsch, Haan, Germany). Samples were aseptically transferred to bead-beating tubes with two sterilised tungsten grinding balls and kept at -20°C until grinding. Samples were ground at 30 hertz in five-minute intervals for up to 20 minutes. Samples were kept frozen by placing them on ice between grindings. Genomic DNA from ground bark samples was then extracted using the SYNERGY 2.0 Plant DNA extraction kit (OPS Diagnostics LLC, US), according to the manufacturer’s instruction, but with doubled amounts of sample and extraction buffer. Genomic DNA from soil samples (0.5 g) was extracted using the FASTDNA Spin Kit (MP Bio). Metagenomic sequencing was conducted at Hudson Genomics. Shotgun metagenomic libraries were prepared using Tecan Celero WGS Libraries and sequenced on the NextSeq2000 platform (2 × 100 bp).

### Quantitative PCR

Quantitative PCR targeting the bacterial-specific 16S rRNA gene V5-7 region was performed to estimate bacterial cell count for the subtropical NSW (2021) and temperate Victoria (2023) samples. The primer pairs 799F (5’-AACMGGATTAGATACCCKG-3’) and 1391R (5’-GACGGGCGGTGWGTRCA-3’) were used to minimize amplification of plastid and mitochondrial DNA^48^. Each 20 µl PCR reaction contained 1 µl of the purified DNA sample (as template), 10 µl of 2 × LightCycler 480 SYBR Green I Master (Millennium Science), and a pair of primers at 0.4 µM each. The PCR was carried out using by the following programs; 3 min at 95°C, and then 50 cycles of 30 sec at 95°C, 30 sec at 53°C, and 30 sec at 72°C for data collection in the Applied Biosystems QuantStudio 7 (ThermoFisher Scientific). The V5-7 region amplicon standards were prepared by amplifying *Escherichia coli* gDNA using the same setting and purified the amplicon using Monarch DNA Gel Extraction Kit (New England BioLabs). Unspecific binding and amplification were tracked using the melt curve which initiated at the end of the PCR stage by the following programs; 15 sec at 95°C, 1 min at 60°C, and 15 sec at 95°C for data collection.

### Metagenome processing, assembly, and binning

Raw metagenomic reads were quality processed through the BBDuk function of BBMap v39.31 (https://sourceforge.net/projects/bbmap/) to remove sequencing adapters (k-mer size: 23, hamming distance: 1), PhiX spike-in (k-mer size: 31, hamming distance: 1), bases from 3’ ends with a Phred score below 20, and resultant reads with lengths shorter than 50 bp. Reads were subsequently mapped to genomes of human, 43 common bacterial contaminants, and 76 tree species (genome accession summarized in **Table S6**) using the BBMap function (maxindel=3 minid=0.95 minhits=2 bwr=0.16 bw=12 untrim quickmatch fast). Mapped reads were removed and summarized in **Table S6** as per recent recommendations^49^.

To recover metagenome-assembled genomes (MAGs), bark and soil metagenomes from the 2022 sampling campaign were assembled individually using metaSPAdes v3.15.3^50^ and collectively (using metagenomes from same tree species) using MEGAHIT v1.2.9^51^, both with the options min k: 27, max k: 127, and k step: 10. Contigs with lengths below 500 bp were discarded. Quality processed short reads were mapped to the assembled contigs using CoverM v0.6.1^52^ with default parameters to generate contig coverage profiles. Genomic binning was performed on contigs with a minimum length of 2000 bp using CONCOCT v1.1.0^53^, MaxBin2 v2.2.7^54^, and MetaBAT2 v2.15^55^. Resulting bins from the same assembly were then dereplicated using DAS_Tool v1.1.6^56^. Applying a common species-level threshold average nucleotide identity of 95%, bins from different assemblies were consolidated to a non-redundant set of MAGs using dRep v3.4.3^57^. A total of 145 new MAGs (95 from bark, 39 from water, 11 from soils) with a minimum completeness of 50% and a maximum contamination of 10%, assessed by CheckM2 v1.0.2^58^, were recovered. MAG taxonomy was assigned by GTDB-Tk v2.3.2^59^ with reference to the Genome Taxonomy Database (GTDB) R08-RS214^60^. A *de novo* phylogenomic tree was further generated using the GTDB bac120 marker set for the 51 diazotrophic MAGs^59^. Marker proteins were identified, aligned, and masked with GTDB-Tk v2.4.1^59^ using default settings, producing a final alignment of 5,035 positions. A maximum-likelihood tree was constructed with IQ-TREE v2.4.0 using the LG+F+R10 model and 1,000 ultrafast bootstraps^61^. The resulting tree was visualized using iTOL v6^62^.

### Metabolic annotations

Gene-centric metagenomic analysis was performed to quantify diazotroph abundance in bark and soil microbial communities. Quality-filtered metagenomic reads over 120 bp were searched against the curated database of *nifH* gene (https://github.com/GreeningLab/GreeningLab-database/tree/main) using DIAMOND v.2.1.14^63^ (query cover > 80%, identity > 50%). Read counts for *nifH* were normalized to reads per kilobase per million (RPKM) by dividing the actual read count by the total number of reads (in millions) and gene length (in kilobases). Reads were also screened for the 13 universal microbial single copy ribosomal marker genes (*rplK*, *rplN*, *rplP*, *rplB*, *rplC*, *rplE*, *rplF*, *rpsJ*, S15P_S13e, *rpsS*, *rpsB*, S5, S7) in PhyloSift^64^ by DIAMOND (query cover > 80%, bitscore > 40) and normalized as above. The average gene copy number of *nifH* in the community was then estimated by dividing RPKM value of the gene by the geomean of RPKM value of the universal single copy ribosomal marker genes. To evaluate if diazotrophs similarly inhabit bark of non-Australian tree species, we reanalyzed published metagenomes of English Oak stem^43^ (*Quercus robur*; from an urban forest in United Kingdom), Norway Spruce bark^19^ (*Picea abies*; from a boreal forest in Sweden), and Hass Avocado bark and soils^44^ (*Persea americana*; from an orchard in Mexico) (Summarized in **Table S6**). These metagenomes were quality processed and surveyed for the *nifH* gene as above.

To identify and profile the distribution and metabolic capability of diazotrophs, the 145 newly recovered MAGs and 114 previously reported bark-associated MAGs were metabolically annotated using DRAM v1.2.4^65^ with the Carbohydrate-Active enZYmes (CAZy) in dbCAN2 database v10^66^ and KEGG protein database^67^ (accessed 22 November 2021). Pathway completeness of nitrogenase catalytic subunits, maturation and accessory proteins, major aerobic and anaerobic respiratory processes, central carbon metabolism, lithotrophy, fermentation, and volatile compound assimilation and oxidation for all MAGs was reported in **Table S7**. MAGs were additionally annotated against the Greening lab metabolic marker gene database (https://github.com/GreeningLab/GreeningLab-database/tree/main) using DIAMOND to identify genes mediating climate-active gas, nitrogen, and sulfur cycling, as described in Leung & Jeffrey et al. 2025^20^.

### Nitrogenase phylogenetic, synteny, and evolutionary analysis

To construct NifH phylogenetic tree, amino acid sequences were aligned using MUSCLE v3.8.1551^68^ (default settings) and further trimmed using TrimAl v1.4.1^69^. Maximum-likelihood trees were constructed using IQ-TREE v2.2.0.3^61^, with best-fit models and 1000 ultrafast bootstraps. All trees were visualized by using iTOL v6^62^. To infer the genetic organisation of nitrogenase, MAGs with long *nifH*-containing contigs were annotated using DRAM v1.3.5^65^ against KEGG, Pfam, MEROPS, and dbCAN databases. The gene clusters were then visualized using the R package gggenes (https://github.com/wilkox/gggenes).

All metagenomic filtered reads from each sample were mapped to an indexed database of the NifH-containing genomes using Bowtie2^70^ (v2.3.5.1; default parameters). The nucleotide diversity and pN/pS ratio of nitrogenase genes (*nifH*, *nifK*, *nifD*) were calculated from these mappings using the profile module of the inStrain program^71^ (v1.9.0, default parameters) at the gene level.

### Nitrogen fixation assays

Bark samples for nitrogen fixation assays were collected from the same trees (northern NSW) on May 8, 2024, stored in the dark and refrigerated overnight prior to incubation. To measure nitrogen fixation rates, the freshly collected bark from each tree **(Table S1)** was dissected into 30 × 5 mm strips, each weighing between 1.0 to 1.5 mg. Triplicate bark strips of each tree and species were carefully placed into sterilised 12 mL glass vials (Exetainer, UK), ensuring integrity of the inner and outer bark natural structures were preserved. For each tree species, three additional vials containing bark were oven-dried at 60°C for four days without prior incubation to determine the background δ^15^N (‰). For heat-killed controls, three bark pieces were randomly selected across different trees and species and autoclaved (121°C, 60 minutes). All treatment and control vials were then capped, the headspace evacuated to remove ambient air, and immediately inoculated with ^15^N-N_2_ gas. To remove any ^15^N compound impurities (e.g., ^15^NH_4_, ^15^NO_x_, ^15^NO, ^15^N_2_O), the ^15^N-N_2_ gas was pre-scrubbed by bubbling it through a 1 M H_2_SO_4_ solution (Dabundo et al., 2014). The inoculated vials were foil wrapped and incubated in the dark at 25°C for four days (ZWY-240 Incubator shaker, Labwit). Finally, the vials were de-capped and oven dried at 60°C for four days. The ^15^N_2_ gas used in the assay was >98% pure (Cambridge Isotopes, USA, Lot I-28469) and was manufactured from ^15^NH_4_Cl (Sigma-Aldrich) according to (Diocares et al., 2006).

To assess the role of methanotrophs in the nitrogen fixation, an additional set of incubations was performed using bark collected from the lower stem (0.5 m) of a *M. quinquenervia* individual from the same subtropical wetland. Bark samples were collected and prepared as described above, and incubated in triplicate under three different gas treatments: (i) stem gas mixture (80% stem gas, 20% ^15^N_2_ gas), (ii) stem gas with methanotrophs inhibitor (78% stem gas, 20% ^15^N_2_, 2% difluoromethane [DFM]) or (iii) stem gas with methane addition (stem gas, 20% ^15^N_2_, 5% of 51,300 ppmv CH_4_). For the latter, given the measured CH_4_ concentration in the stem gas was 655 ppmv, the final CH_4_ concentration in the vials was ∼3,060 ppm (∼0.31% CH_4_). All treatments were incubated for one and two days. Triplicate bark pieces were oven-dried at 60°C without incubation to determine the natural δ^15^N isotopic composition and triplicate autoclaved bark samples, incubated with the stem gas treatment for one and two days, served as heat-killed controls.

### Stem gas collection

Stem gas was collected using a semi-rigid chamber^72^ (40 × 30 cm) fixed at 0.5 m stem height and connected in a closed loop to a desiccant trap (Drierite), air pump (T2-05, Parker Pumps), and 1 L twin-valve foil gas bag (Cali-5-Bond) using gas tubing (Bev-A-Line-IV). One litter of ambient air was used to initially fill the gas bag, and then the air was circulated (0.24 L min^-1^) for 24 hours to allow stem gases to reach equilibrium. Two 5 mL subsamples of stem gas were analysed for CH_4_ and CO_2_ using a small sample induction module (SSIM AO314, Picarro) connected to a factory calibrated cavity ring-down spectroscopy analyzer (CRDS G2201-i, Picarro).

### Measurement of isotopic composition and N fixation rates

The dried bark pieces were chopped and placed into 2 mL Eppendorf tubes containing sterile 5 mm stainless beads (Qiagen). Samples were homogenised into fine powder using a TissueLyser II (Qiagen) at 30 Hz for one minute, with up to four repetitions as needed to ensure complete homogenisation. The δ^15^N of bark samples were run in an elemental analyser (Thermo Fisher Flash EA) coupled to an isotope ratio mass spectrometer (IRMS, Thermo Fisher Delta V plus) via an interface (Thermo Fisher Conflo IV) as described in Carvalho et al. (2024)^73^. Briefly, approximately 15 ± 0.6 mg of homogenised sample was placed into a capsule and the net weight was measured using a microbalance (sensitivity of ± 1 µg, Sartorius). Given the high carbon content in bark, a second GC column (Mol Sieve 5A plot Column) was used at 40°C to trap CO_2_ and separate CO from N_2_. Every 16 samples, the reactor was cleaned and the temperature of the Mol Sieve 5A GC column was raised to 300°C to release accumulated CO_2_. Samples were measured along working standards (glycine, δ^15^N = 2.0‰; collagen, δ^15^N = 4.8‰), ensuring precision of δ^15^N within ± 0.3‰. Nitrogen fixation rates were calculated using well-established methods^74^ and upscaled to 4 m height based on species-specific parameters such as bark thickness (mm) and density (g/cm^2^) and bark surface area (m^2^). When δ^15^N background values were higher than some of the ^15^N-N_2_ incubated samples, the lowest was selected as the δ^15^N baseline. Because of the variability in the natural abundance δ^15^N between trees of the same species, BNF rates were calculated based on a specific δ^15^N baseline for each tree.

### Biogeochemical analyses and ancillary data

Paired bark and soil samples from each species were analysed for total N and C content (%), macronutrient and micronutrient composition (P, Mo, V, Fe, Ni, Cu; mg kg^-1^) at Environmental Analysis Laboratory (EAL, Southern Cross University) **(Table S10)**. Additionally, soil, leaf litter and leaves from each tree were collected on October 15, 2025, dried at 60°C for 3 days, ground to a fine powder and analysed for total C and N (%) and stable nitrogen isotope ratios (δ^15^N, ‰) using an EA-IRMS as described above **(Tables S2-5).** Additional forest parameters were recorded at each sampling site, including tree density (trees ha^-1^), tree height (m), diameter at breast height (DBH, cm), bark thickness (mm) and calculated stem surface area (m^2^). Adjacent soil volumetric water content (VWC %) was measured using a soil moisture probe. Surface water depth was recorded where present in wetland sites. Complete metadata for the sampled trees in northern NSW are provided in **Table S1**.

### Statistical analyses

All statistical analyses were performed in R version 4.5.0. The nitrogen stable isotope signatures (δ^15^N) in the bark samples of the eight tree species were analysed using one-way analysis of variance (ANOVA). Normality and homoscedasticity of the data were assessed using the Shapiro-Wilk (W = 0.95, *p* = 0.21) and Levene’s tests (F = 0.57, *p* = 0.77), respectively. Post-hoc pairwise comparisons with Tukey adjustment were conducted using estimated marginal means (EMMs) via the *emmeans* R package^75^. Differences in total carbon (%C), total nitrogen (%N), and carbon-to-nitrogen ratio (C/N) and δ^15^N across forest compartments (bark, leaves, leaf litter and soil) were assessed using the non-parametric Kruskal-Wallis test, due to violation of normality assumptions. Pairwise comparisons were performed by post-hoc Dunn’s test (Bonferroni adjustment). Significance was set at α = 0.05 for all tests.

Nitrogen fixation rates (ng N g^-1^ day^-1^) across species were analysed using multiple modelling approaches to account for the nested structure of the data and the non-normal distribution of the response variable. Candidate models included analysis of variance (ANOVA) with Box-Cox transformation and tree as fixed effect, linear mixed-effects models (LME) with Box-Cox transformation and tree as random effect, generalised linear models (GLM) with Gamma distribution and log link without and with random effects (GLMM) and GLM with link log Gamma distribution and tree nested within species as fixed effects **(Table S11)**. Box-Cox transformation was performed using the boxcox function (*MASS* package^76^) to determine the optimal power transformation (λ) for normalising the data. All candidate models included ‘species’ as a fixed effect and ‘tree identity’ either as random effect (LME, GLMM) or fixed effect (ANOVA, nested GLM) to account for potential intra-species variability.

Candidate models were evaluated using Akaike Information Criterion (AIC), Bayesian Information Criterion (BIC), log likelihood values, and residual diagnostics including residuals vs. fitted plots, Q-Q plots, Shapiro-Wilk test for normality, and Levene’s test for homogeneity of variances. The LME model with Box-Cox transformation (λ = 0.222) provided substantially better fit (AIC = 268.35, BIC = 292.43) compared to all alternative models (**Table S11**). The LME was implemented using the lme function in the *nlme* R package^77^ with tree identity as a random intercept. While GLMM with Gamma distribution and log link offered biological interpretability on the original scale, it was not selected due to poorer fit (AIC = 455.02; **Table S11**). Model assumptions were evaluated by inspecting patterns in residuals versus fitted values, quantile-quantile (QQ) plots, Shapiro-Wilk tests for normality (W = 0.98, *p* = 0.35) and Levene’s test to confirm homogeneity of variances (F = 0.37, *p* = 0.93).

The significance of fixed effects (species) was assessed using Type II Wald tests via the *car* package^78^. Post-hoc pairwise comparisons between species were performed using EMMs with Benjamini-Hochberg adjustment to control false discovery rate. Results were back-transformed to the original scale (ng N g^-1^ day^-1^) using inverse Box-Cox transformation for biological interpretation, while p-values were obtained from the transformed scale where model assumptions were satisfied.

Nitrogen fixation rates measured in *M. quinquenervia* during the CH_4_/DFM assays were analysed using a two-way ANOVA with fixed effects to assess the effect of treatment (stem gas, stem gas + CH_4_, stem gas + DFM) and incubation time. Tukey’s HSD post-hoc test was performed for pairwise comparisons. Normality and homoscedasticity of the data were assessed using the Shapiro-Wilk (W = 0.90, *p* = 0.06) and Levene’s tests (F = 0.75, *p* = 0.60), respectively.

## Supporting information

Table S6. Tree-fix metagenome info & Gene abundance

Table S7. Tree-fix MAG summary

## Acknowledgements

This work was supported by the Australian Research Council, namely Discovery Project grants (DP210100096 to D.T.M, S.G.J; DP210101595 to C.G., P.M.L.C., W.W.W.), Discovery Early Career Research Award fellowships (DE240100338 to L.C.J; DE250101210 to P.M.L.), and a Future Fellowship (FT240100502 to C.G.), as well as awards from the Hermon Slade Foundation (to L.C.J., D.T.M., S.G.J, C.G.) and Holsworth Wildlife Foundation (to P.M.L.). We acknowledge Tweed Shire Council, NSW NPWS for sampling permissions (permit SL102622) and the Parks and Wildlife Commission, Northern Territory (permit 75926). We thank Thanavit Jirapanjawat, Clément Duvert, Francesco Ulloa Cedamanos, Kaline de Mello, Lindsay Hutley, Alan Anderson, Naomi Jeffrey, Matheus Carvalho de Carvalho, Tiffeny Byrnes, Iain Alexander, and Matt Veness for field and laboratory assistance and Sean Bay for conducting preliminary MAG analysis.

## Author contributions

C.G., D.T.M., P.M.L., L.C.J., and S.G.J. conceived this study. D.T.M., C.G., P.G.A., L.C.J., S.G.J., and P.M.L. designed this study. P.G.A., L.C.J., J.D., S.C., S.E., D.T.M., M.H, B.H. and P.M.L. conducted fieldwork. P.M.L., C.Z., C.G., A.J.K., and X.D. conducted metagenomic analyses. C.Z., X.D., and C.G. conducted nitrogenase phylogenetic and synteny analysis. P.G.A., D.T.M., L.C.J., D.E., P.L.M.C, B.H, and W.W.W. conducted nitrogen fixation assays. P.G.A. performed statistical analysis. C.G., P.G.A., D.T.M., C.Z., L.C.J, X.D., A.J.K., and P.M.L. wrote and illustrated the paper with input from all authors.

## Data availability

The metagenomes and metagenome-assembled genomes sequenced in this study are being deposited in NCBI.

