## Supplementary Information for "Bark-associated diazotroph communities are a cryptic source of nitrogen in forests"

**Figure S1. Map showing bark and sampling locations used for nitrogen fixation rate estimates and metagenomic sequencing.** Locality maps showing the different forest biomes where bark was collected from for the ^15^N_2_ labelling assays and metagenomics from (sub)tropical and temperate Australian forests. World map shows location of other published bark metagenomes used in this study.

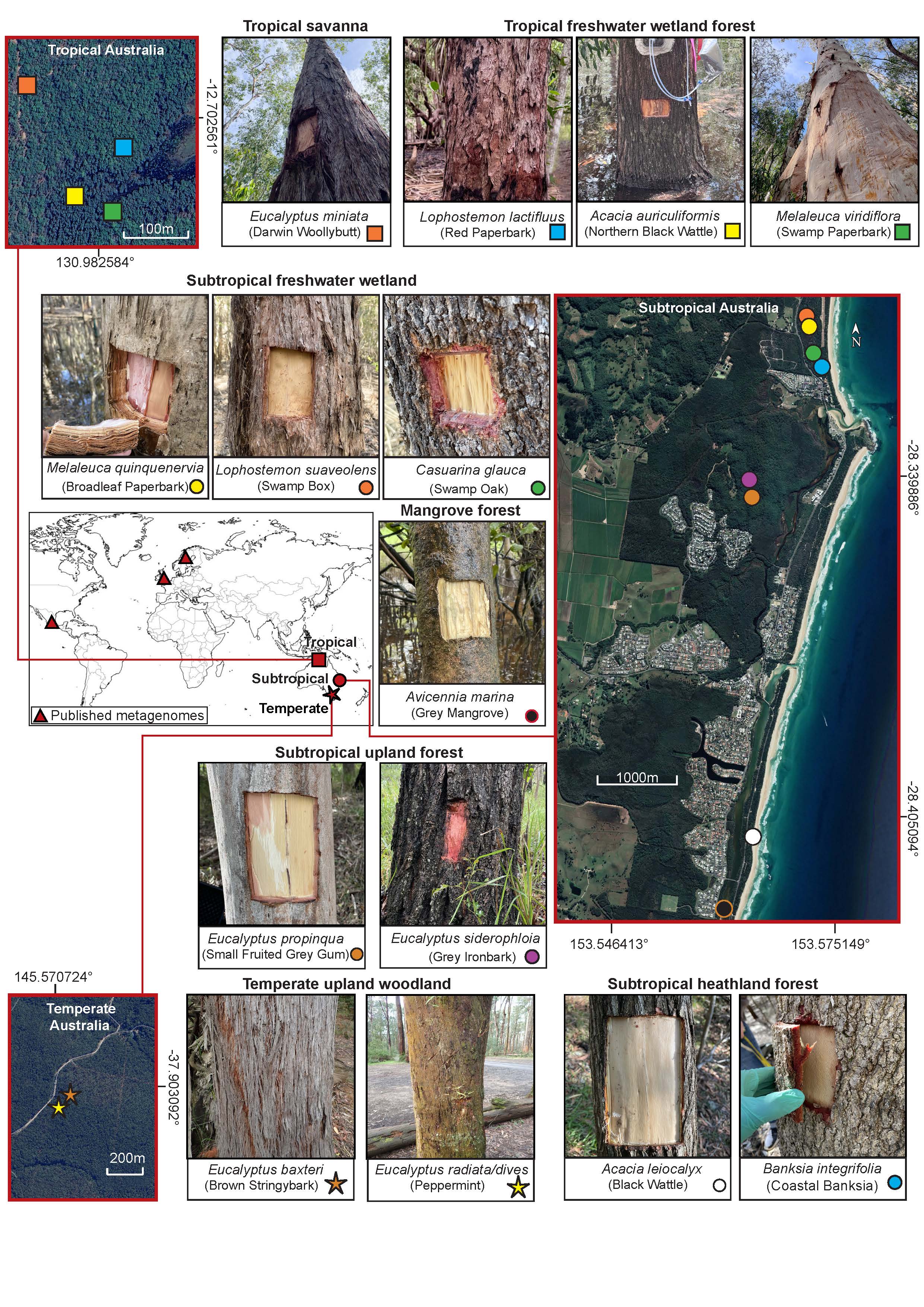

**Figure S2. Correlation of diazotroph abundance and nitrogen fixation signatures in tree bark.** A linear regression analysis shows the extent to which bark diazotroph community abundance (based on proportion of cells encoding *nifH*) are correlated with bark δ^15^N signatures.

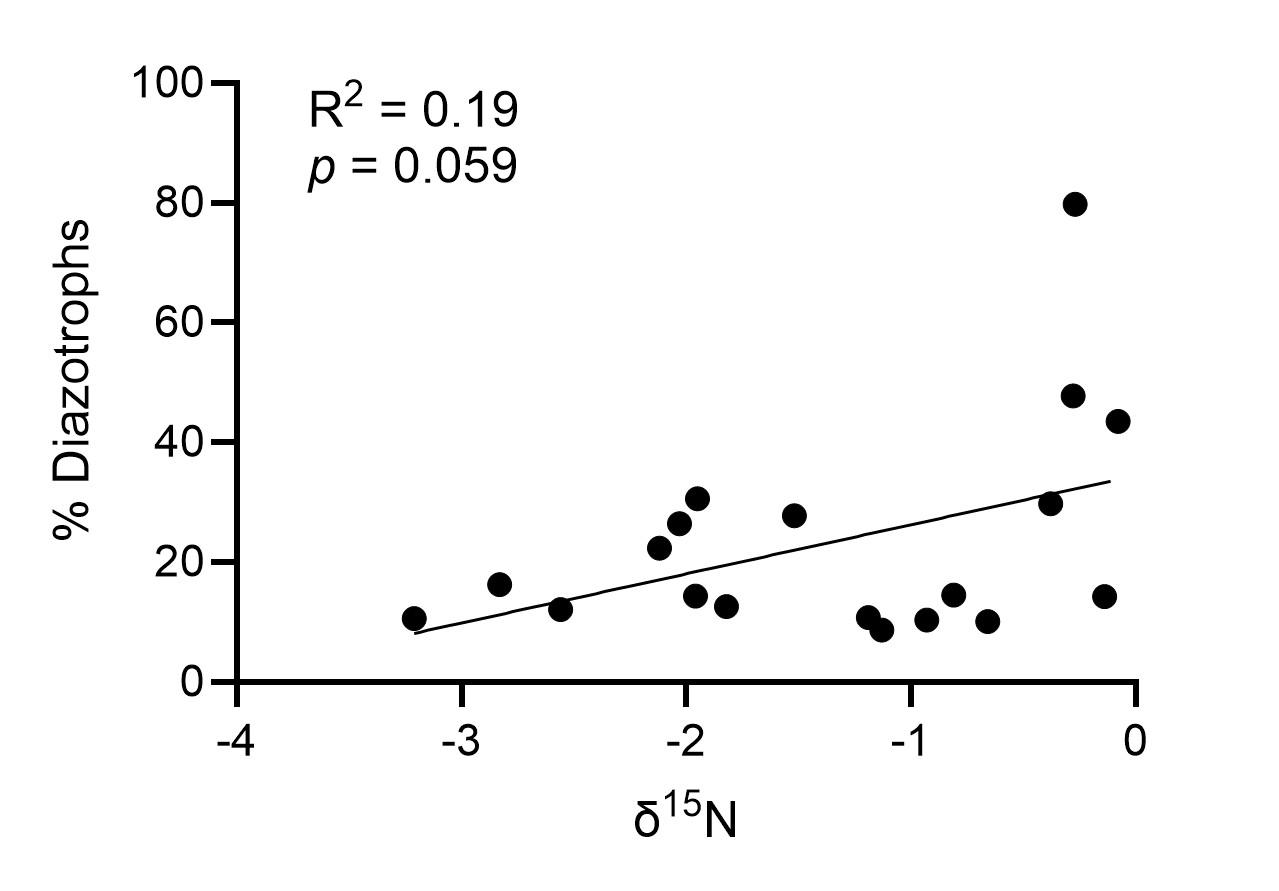

**Figure S3. Phylogenetic diversity of the nitrogenases encoded by tree microbial communities.** The tree shows reference sequences from nitrogenase lineages (Group I, II, III, IV-A, and VII) and nitrogenase-like proteins in black. Unbinned contigs from tree bark are shown in red and adjacent soils / waters are shown in blue.

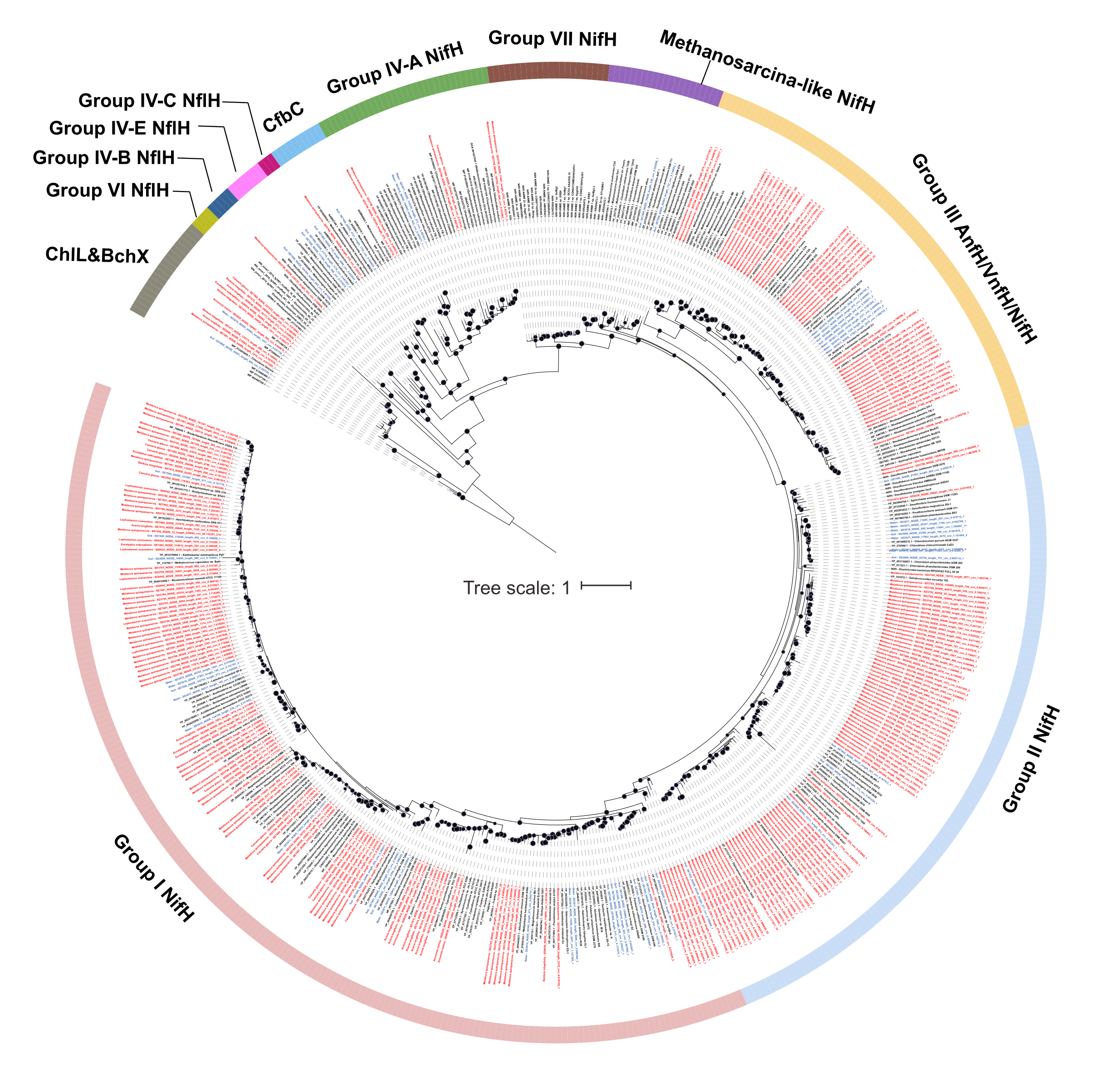

**Figure S4. Nitrogen fixation rates among tree species and individual trees across forest types.** N fixation rates (ng N g^-1^ day^-1^) measured in the ^15^N_2_ labelling assays for individual trees of different species across four forest ecosystems: freshwater wetland, heathland, upland and mangrove forest. Box plots indicate median (central line), interquartile range (box), and 1.5× interquartile range (whiskers). Wetland forests exhibited the highest variation in N fixation rates, particularly in *Melaleuca quinquenervia*, while upland forests showed the lowest overall rates. Intraspecific variation among individual trees was substantial in several species, suggesting tree-specific microbiome differences may influence N fixation capacity. Rates vary at the levels of ecosystem type (panels show four ecosystem types), tree species (data grouped by eight tree species), individual tree (coloured boxes show three individual trees per species), and bark subsample (individual points show three replicates per tree, n = 3).

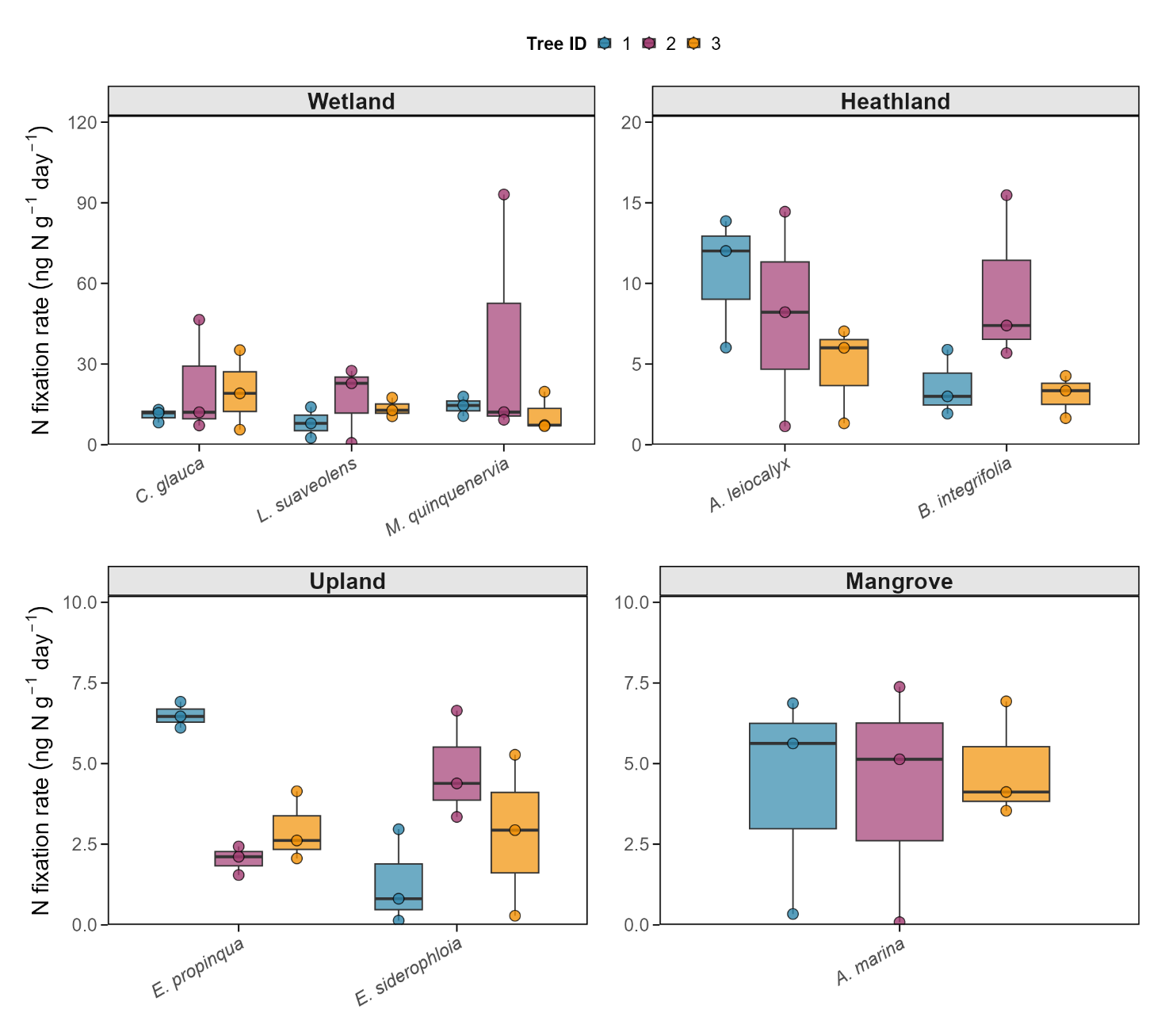

**Figure S5. Comparison of diazotroph abundance based on *nifH* between different sources and tree heights. (a)** Diazotroph proportion in all sequenced bark (n = 72), soil (n = 30) and heartwood (n = 11) from Australian forests. Statistical differences were tested by student’s *t* tests (** *p* < 0.001, * *p* < 0.05, ns *p* > 0.05). **(b)** Diazotroph proportion in outer bark sampled along various tree height for five tree species.

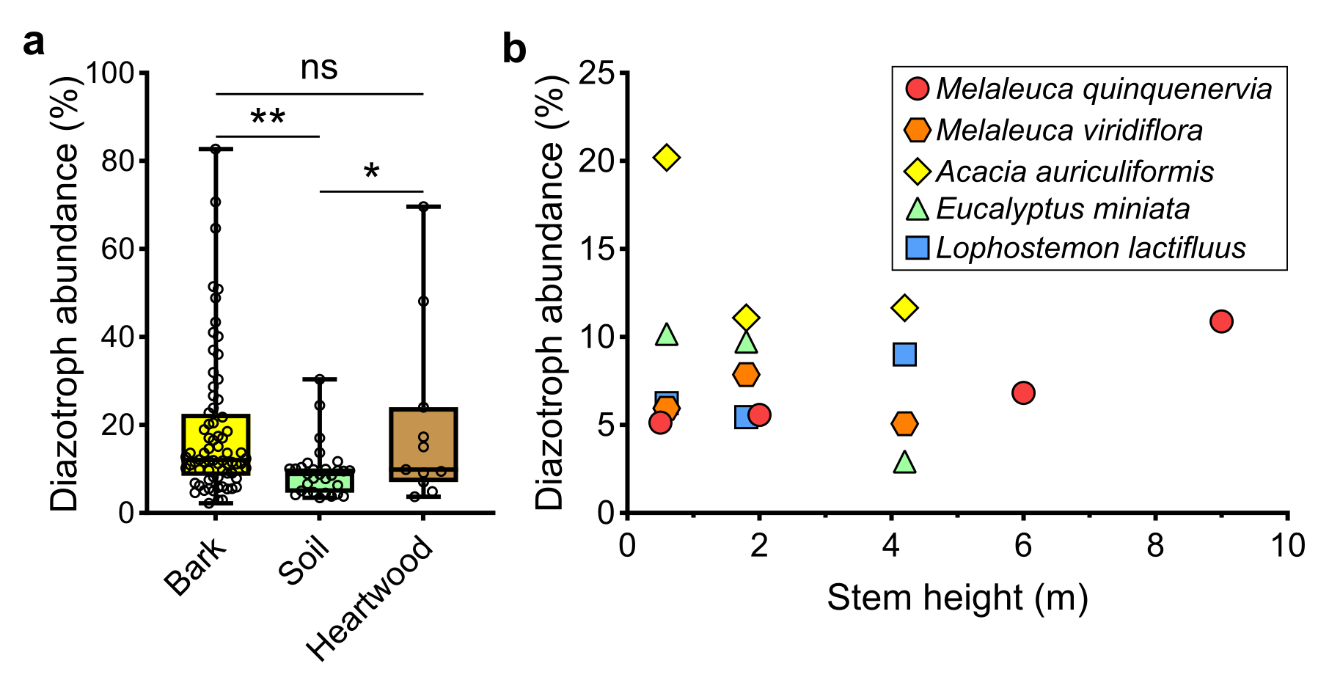

**Table S1. Tree metadata for metagenomic and biogeochemical analysis**. Characteristics and environmental data for 24 trees (three trees per species) from eight species across mangrove (*Avicennia marina*), freshwater wetland (*Casuarina glauca, Melaleuca quinquenervia, Lophostemon suaveloens*), upland (*Eucalyptus siderophloia*, *Eucalyptus propinqua*) and heathland (*Banksia integrifolia, Acacia leiocalyx*) ecosystems in northern New South Wales, Australia. Data include location, elevation, stem dimensions, bark properties, moisture content, and soil/water conditions.

| **Tree species** | **Common name** | **Tree ID** | **Lat., Long.** | **Elevation (m)** | **Sampling height (cm)** | **Stem circum. (cm)** | **Bark thickness (mm)** | **Bark moisture (%)** | **DBH (cm)** | **Tree height (m)** | **Wood core length (mm)** | **Wood core moisture (%)** | **Soil moisture (%)** | **Surface water depth (cm)** |
| --- | --- | --- | --- | --- | --- | --- | --- | --- | --- | --- | --- | --- | --- | --- |
| *Avicennia marina* | Grey Mangrove | 1 | -28.41271°, 153.56123° | 2.7 | 84 | 36 | 4 | 56.3 | 13.5 | 9.9 | 35 | 39.7 | 100 | - |
|  |  | 2 |  | -2.7 | 97 | 30.4 | 10 | 45.3 | 9.6 | 10.2 | **40 |  | 100 | - |
|  |  | 3 |  | -2.7 | 112 | 28 | 9 | 45.4 | 9.1 | 10.5 | 38 |  | 100 | - |
| *Acacia leiocalyx* | Black wattle | 1 | -28.40520°, 153.56360° | 22.3 | 104 | 54.2 | 8 | 48.6 | 17 | 15.8 | **18 | 40.8 | 8.4 | - |
|  |  | 2 |  | 22.5 | 105 | 51.9 | 11 | 44.6 | 14.8 | 13.8 | 27 |  | 11.1 | - |
|  |  | 3* |  | 14.8 | 87 | 37.8 | 7 | 44.3 | 11.3 | 14.1 | 29 |  | 9.7 | - |
| *Banksia integrifolia* | Coastal Banksia | 1 | -28.35173°, 153.57227° | 21.7 | 86.5 | 123.4 | 15 | 56.4 | 42 | 16.4 | *50 | 56 | 18.4 | - |
|  |  | 2 |  | 21.7 | 77 | 120 | 14 | 55.2 | 28.2 | 9.4 | 49 |  | 11.8 | - |
|  |  | 3 |  | 25.9 | 84.5 | 143 | 15 | 53.1 | 43.5 | 12.2 | 52 |  | 12.4 | - |
| *Casuarina glauca* | Swamp Oak | 1 | -28.35187°, 153.57179° | 5.1 | 35 | 114 | 19 | 44.8 | 31.5 | 17.5 | **48 | 34.8 | 34.8 | 45 |
|  |  | 2 |  | 3.6 | 65 | 76 | 15 | 43.9 | 25 | 18.5 | 47 |  | 34.8 | 3 |
|  |  | 3 |  | 4.6 | 60 | 75 | 13 | 45.5 | 22.5 | 15.3 | 50 |  | 100 | 91 |
| *Eucalyptus propinqua* | Small Fruited Grey Gum | 1 | -28.36964°, 153.56358° | 51.4 | 85 | 162.5 | 11 | 49.6 | 50.8 | 22 | **38 | 34.5 | 21.3 | - |
|  |  | 2 |  | 44.2 | 107 | 71.8 | 5 | 54.8 | 22.2 | 18.8 | 34 |  | 13.1 | - |
|  |  | 3 |  | 42.7 | 109.5 | 64.9 | 20 | 52.2 | 20.2 | 19.6 | 28 |  | 19.6 | - |
| *Eucalyptus siderophloia* | Iron Bark | 1 | -28.36996°, 153.57121° | 58.2 | 84.5 | 133.8 | 29 | 34.3 | 43.8 | 24.4 | **35 | 27.1 | 19.9 | - |
|  |  | 2 |  | 69.8 | 78 | 164.5 | 18 | 35.2 | 52.4 | 21.3 | 40 |  | 19.9 | - |
|  |  | 3 |  | 63.7 | 104 | 87 | 11 | 33.5 | 27 | 18.2 | 45 |  | 23.8 | - |
| *Lophostemon suaveolens* | Swamp Box | 1 | -28.34924°, 153.57121° | 25.1 | 108 | 76.3 | 15 | 41.0 | 25 | 11.8 | **41 | 43.7 | 25.8 | - |
|  |  | 2 |  | 25.2 | 81.1 | 108.9 | 11 | 39.6 | 34.8 | 12.4 | 46 |  | 6.5 | - |
|  |  | 3 |  | 43.3 | 86 | 30.1 | 15 | 46.7 | 29.4 | 15.1 | 49 |  | 100 | 23 |
| *Melaleuca quinquenervia* | Paperbark | 1* | -28.34839°, 153.57107° | 25.5 | 61 | 89 | 22 | 66.4 | 28.3 | 14.4 | **46 | 48.1 | 44.1 | - |
|  |  | 2* |  | 33.2 | 45 | 120 | 32 | 69.6 | 35.7 | 14.5 | 39 |  | 37.3 | - |
|  |  | 3* |  | 34.4 | 62 | 100 | 24 | 64.5 | 31.8 | 14.8 | 54 |  | 41.0 | - |

* Indicates trees sampled in 2024 for ^15^N_2_-labelling assays different from the sampled trees for metagenomics in 2021, ** indicates wood cores sequenced.

**Table S2. Carbon and nitrogen content and stable isotope signatures in bark.** Elemental composition (%N, %C), stable isotope ratios (δ^15^N, ‰) and carbon-to-nitrogen ratios (C/N) measured in bark of eight Australian tree species by EA-IRMS. Data represent measurements from three different trees per species (Tree ID).

| **Species** | **Common name** | **Tree ID** | **Sample type** | **%N** | **%C** | **δ^15^N** | **C/N** |
| --- | --- | --- | --- | --- | --- | --- | --- |
| *Acacia leiocalyx* | Black wattle | 1 | Bark | 1.06 | 48.30 | -1.95 | 45.41 |
| *Acacia leiocalyx* | Black wattle | 2 | Bark | 0.67 | 48.30 | -2.03 | 71.81 |
| *Acacia leiocalyx* | Black wattle | 3 | Bark | 0.73 | 48.30 | -1.52 | 66.02 |
| *Eucalyptus siderophloia* | Iron bark | 1 | Bark | 0.27 | 53.00 | -1.96 | 195.06 |
| *Eucalyptus siderophloia* | Iron bark | 2 | Bark | 0.15 | 53.00 | -3.21 | 364.65 |
| *Eucalyptus siderophloia* | Iron bark | 3 | Bark | 0.19 | 53.00 | -2.83 | 273.15 |
| *Avicennia marina* | Grey mangrove | 1 | Bark | 0.71 | 48.00 | 4.47 | 67.66 |
| *Avicennia marina* | Grey mangrove | 2 | Bark | 0.33 | 48.00 | 4.52 | 144.60 |
| *Avicennia marina* | Grey mangrove | 3 | Bark | 0.47 | 48.00 | 4.44 | 101.11 |
| *Eucalyptus propinqua* | Small fruited grey gum | 1 | Bark | 0.17 | 43.30 | -2.56 | 259.75 |
| *Eucalyptus propinqua* | Small fruited grey gum | 2 | Bark | 0.19 | 43.30 | -0.93 | 227.30 |
| *Eucalyptus propinqua* | Small fruited grey gum | 3 | Bark | 0.22 | 43.30 | -2.12 | 198.57 |
| *Banksia integrifolia* | Banksia | 1 | Bark | 0.20 | 51.10 | -0.14 | 252.44 |
| *Banksia integrifolia* | Banksia | 2 | Bark | 0.20 | 51.10 | -1.82 | 249.50 |
| *Banksia integrifolia* | Banksia | 3 | Bark | 0.22 | 51.10 | -1.19 | 234.86 |
| *Lophostemon suaveolens* | Swamp box | 1 | Bark | 0.35 | 49.90 | -0.66 | 142.86 |
| *Lophostemon suaveolens* | Swamp box | 2 | Bark | 0.31 | 49.90 | -0.81 | 158.82 |
| *Lophostemon suaveolens* | Swamp box | 3 | Bark | 0.34 | 49.90 | 0.51 | 145.37 |
| *Melaleuca quinquenervia* | Paperbark | 1 | Bark | 0.45 | 58.50 | 0.18 | 130.99 |
| *Melaleuca quinquenervia* | Paperbark | 2 | Bark | 0.20 | 58.50 | -0.38 | 290.63 |
| *Melaleuca quinquenervia* | Paperbark | 3 | Bark | 0.37 | 58.50 | -0.28 | 158.61 |
| *Casuarina glauca* | Swamp oak | 1 | Bark | 0.49 | 46.70 | -0.27 | 94.37 |
| *Casuarina glauca* | Swamp oak | 2 | Bark | 0.53 | 46.70 | -1.13 | 88.94 |
| *Casuarina glauca* | Swamp oak | 3 | Bark | 0.46 | 46.70 | -0.08 | 102.52 |

**Table S3. Carbon and nitrogen content and stable isotope signatures in soil.** Elemental composition (%N, %C), stable isotope ratios (δ^15^N, ‰) and carbon-to-nitrogen ratios (C/N) for the underlying soil of eight Australian tree species measured by EA-IRMS. Data represent measurements from three different trees per species (Tree ID).

| **Species** | **Common name** | **Tree ID** | **Sample type** | **%N** | **%C** | **δ^15^N** | **δ^13^C** | **CN** |
| --- | --- | --- | --- | --- | --- | --- | --- | --- |
| *Acacia leiocalyx* | Black wattle | 1 | Soil | 0.12 | 2.29 | 1.00 | -27.0 | 18.35 |
| *Acacia leiocalyx* | Black wattle | 2 | Soil | 0.17 | 3.22 | 0.86 | -27.6 | 19.04 |
| *Acacia leiocalyx* | Black wattle | 3 | Soil | 0.09 | 1.51 | 0.77 | -27.5 | 17.10 |
| *Eucalyptus siderophloia* | Iron bark | 1 | Soil | 0.34 | 9.60 | 0.51 | -28.0 | 28.41 |
| *Eucalyptus siderophloia* | Iron bark | 2 | Soil | 0.29 | 5.64 | 1.16 | -28.5 | 19.74 |
| *Eucalyptus siderophloia* | Iron bark | 3 | Soil | 0.18 | 4.64 | 1.38 | -27.2 | 25.54 |
| *Avicennia marina* | Grey mangrove | 1 | Soil | 0.56 | 8.87 | 4.38 | -26.2 | 15.70 |
| *Avicennia marina* | Grey mangrove | 2 | Soil | 0.09 | 1.49 | 4.02 | -27.2 | 17.45 |
| *Avicennia marina* | Grey mangrove | 3 | Soil | 0.40 | 5.98 | 4.01 | -25.8 | 15.12 |
| *Eucalyptus propinqua* | Small fruited grey gum | 1 | Soil | 0.28 | 5.18 | 2.11 | -25.8 | 18.70 |
| *Eucalyptus propinqua* | Small fruited grey gum | 2 | Soil | 0.34 | 7.73 | 1.53 | -26.4 | 22.86 |
| *Eucalyptus propinqua* | Small fruited grey gum | 3 | Soil | 0.43 | 10.26 | 0.79 | -28.1 | 23.97 |
| *Banksia integrifolia* | Banksia | 1 | Soil | 0.21 | 5.28 | 0.11 | -30.3 | 24.60 |
| *Banksia integrifolia* | Banksia | 2 | Soil | 0.11 | 2.28 | 1.97 | -26.1 | 21.06 |
| *Banksia integrifolia* | Banksia | 3 | Soil | 0.05 | 1.07 | 2.56 | -27.0 | 20.84 |
| *Lophostemon suaveolens* | Swamp box | 1 | Soil | 0.07 | 1.74 | 2.01 | -28.8 | 25.22 |
| *Lophostemon suaveolens* | Swamp box | 2 | Soil | 1.25 | 28.19 | 3.23 | -28.7 | 22.53 |
| *Lophostemon suaveolens* | Swamp box | 3 | Soil | 0.66 | 15.00 | 2.88 | -27.3 | 22.71 |
| *Melaleuca quinquenervia* | Paperbark | 1 | Soil | 0.17 | 5.39 | 1.71 | -29.9 | 30.87 |
| *Melaleuca quinquenervia* | Paperbark | 2 | Soil | 0.48 | 18.42 | 1.39 | -30.4 | 38.09 |
| *Melaleuca quinquenervia* | Paperbark | 3 | Soil | 0.95 | 36.38 | 1.14 | -30.3 | 38.24 |
| *Casuarina glauca* | Swamp oak | 1 | Soil | 0.33 | 6.70 | -0.66 | -29.8 | 20.50 |
| *Casuarina glauca* | Swamp oak | 2 | Soil | 0.57 | 12.34 | -0.29 | -29.3 | 21.78 |
| *Casuarina glauca* | Swamp oak | 3 | Soil | 1.40 | 28.22 | -0.73 | -30.1 | 20.20 |

**Table S4. Carbon and nitrogen content and stable isotope signatures in leaf litter.** Elemental composition (%N, %C), stable isotope ratios (δ^15^N, ‰) and carbon-to-nitrogen ratios (C/N) for the underlying leaf litter of eight Australian tree species measured by EA-IRMS. Data represent measurements from three different trees per species (Tree ID).

| **Species** | **Common name** | **Tree ID** | **Sample type** | **%N** | **%C** | **δ^15^N** | **δ^13^C** | **C/N** |
| --- | --- | --- | --- | --- | --- | --- | --- | --- |
| *Acacia leiocalyx* | Black wattle | 1 | Leaf litter | 1.13 | 53.11 | -0.87 | -31.5 | 47.18 |
| *Acacia leiocalyx* | Black wattle | 2 | Leaf litter | 1.12 | 54.05 | -0.86 | -31.6 | 48.10 |
| *Acacia leiocalyx* | Black wattle | 3 | Leaf litter | 1.26 | 53.40 | 0.06 | -33.1 | 42.35 |
| *Eucalyptus siderophloia* | Iron bark | 1 | Leaf litter | 1.18 | 52.38 | -2.42 | -30.7 | 44.24 |
| *Eucalyptus siderophloia* | Iron bark | 2 | Leaf litter | 1.21 | 50.83 | -2.69 | -30.5 | 42.13 |
| *Eucalyptus siderophloia* | Iron bark | 3 | Leaf litter | 1.11 | 42.23 | -1.74 | -30.4 | 38.04 |
| *Avicennia marina* | Grey mangrove | 1 | Leaf litter | 0.65 | 35.64 | 5.80 | -28.6 | 54.82 |
| *Avicennia marina* | Grey mangrove | 2 | Leaf litter | 0.53 | 44.93 | 5.77 | -26.8 | 85.30 |
| *Avicennia marina* | Grey mangrove | 3 | Leaf litter | 0.57 | 44.00 | 6.57 | -27.9 | 76.80 |
| *Eucalyptus propinqua* | Small fruited grey gum | 1 | Leaf litter | 0.98 | 49.91 | -2.10 | -29.1 | 51.07 |
| *Eucalyptus propinqua* | Small fruited grey gum | 2 | Leaf litter | 0.74 | 29.86 | -1.06 | -28.7 | 40.16 |
| *Eucalyptus propinqua* | Small fruited grey gum | 3 | Leaf litter | 1.23 | 52.22 | -4.20 | -29.8 | 42.32 |
| *Banksia integrifolia* | Banksia | 1 | Leaf litter | 0.52 | 89.61 | -0.11 | -30.5 | 171.98 |
| *Banksia integrifolia* | Banksia | 2 | Leaf litter | 0.53 | 50.99 | -1.41 | -30.5 | 96.84 |
| *Banksia integrifolia* | Banksia | 3 | Leaf litter | 1.05 | 51.82 | -0.97 | -29.9 | 49.21 |
| *Lophostemon suaveolens* | Swamp box | 1 | Leaf litter | 0.82 | 49.22 | 0.20 | -28.8 | 59.78 |
| *Lophostemon suaveolens* | Swamp box | 2 | Leaf litter | 0.97 | 55.82 | 0.55 | -32.5 | 57.33 |
| *Lophostemon suaveolens* | Swamp box | 3 | Leaf litter | 1.09 | 54.84 | 0.04 | -30.3 | 50.43 |
| *Melaleuca quinquenervia* | Paperbark | 1 | Leaf litter | 1.14 | 54.51 | 0.42 | -30.3 | 47.64 |
| *Melaleuca quinquenervia* | Paperbark | 2 | Leaf litter | 0.97 | 57.99 | 0.96 | -31.8 | 59.83 |
| *Melaleuca quinquenervia* | Paperbark | 3 | Leaf litter | 0.90 | 58.22 | 1.87 | -32.3 | 64.60 |
| *Casuarina glauca* | Swamp oak | 1 | Leaf litter | 1.43 | 51.35 | -1.62 | -31.4 | 35.92 |
| *Casuarina glauca* | Swamp oak | 2 | Leaf litter | 1.65 | 52.17 | -2.14 | -31.2 | 31.65 |
| *Casuarina glauca* | Swamp oak | 3 | Leaf litter | 1.70 | 51.64 | -2.04 | -31.3 | 30.41 |

**Table S5. Carbon and nitrogen content and stable isotope signatures in living leaves.** Elemental composition (%N, %C), stable isotope ratios (δ^15^N, ‰) and carbon-to-nitrogen ratios (C/N) for leaves of eight Australian tree species measured by EA-IRMS. Data represent measurements from three different trees per species (Tree ID).

| **Species** | **Common name** | **Tree ID** | **Sample type** | **%N** | **%C** | **δ^15^N** | **δ^13^C** | **C/N** |
| --- | --- | --- | --- | --- | --- | --- | --- | --- |
| *Acacia leiocalyx* | Black wattle | 1 | Leaves | 2.63 | 52.73 | -1.00 | -31.2 | 20.02 |
| *Acacia leiocalyx* | Black wattle | 2 | Leaves | 2.34 | 52.87 | 0.43 | -31.5 | 22.63 |
| *Acacia leiocalyx* | Black wattle | 3 | Leaves | 2.66 | 53.06 | -1.41 | -31.4 | 19.96 |
| *Eucalyptus siderophloia* | Iron bark | 1 | Leaves | 1.24 | 52.96 | -1.96 | -31.7 | 42.67 |
| *Eucalyptus siderophloia* | Iron bark | 2 | Leaves | 0.94 | 50.99 | -2.95 | -30.4 | 54.07 |
| *Eucalyptus siderophloia* | Iron bark | 3 | Leaves | 1.47 | 50.17 | -2.62 | -30.5 | 34.16 |
| *Avicennia marina* | Grey mangrove | 1 | Leaves | 2.22 | 40.52 | 3.95 | -29.2 | 18.26 |
| *Avicennia marina* | Grey mangrove | 2 | Leaves | 1.76 | 43.04 | 4.36 | -29.4 | 24.41 |
| *Avicennia marina* | Grey mangrove | 3 | Leaves | 1.57 | 44.49 | 4.58 | -30.0 | 28.36 |
| *Eucalyptus propinqua* | Small fruited grey gum | 1 | Leaves | 1.35 | 50.49 | -1.25 | -31.2 | 37.35 |
| *Eucalyptus propinqua* | Small fruited grey gum | 2 | Leaves | 1.50 | 52.28 | -1.18 | -29.5 | 34.94 |
| *Eucalyptus propinqua* | Small fruited grey gum | 3 | Leaves | 1.22 | 49.51 | -1.20 | -32.9 | 40.72 |
| *Banksia integrifolia* | Banksia | 1 | Leaves | 0.90 | 48.23 | 7.63 | -32.4 | 53.47 |
| *Banksia integrifolia* | Banksia | 2 | Leaves | 1.06 | 50.19 | -1.72 | -29.7 | 47.45 |
| *Banksia integrifolia* | Banksia | 3 | Leaves | 1.07 | 37.03 | -1.89 | -31.6 | 34.55 |
| *Lophostemon suaveolens* | Swamp box | 1 | Leaves | 1.27 | 51.19 | 0.10 | -31.5 | 40.33 |
| *Lophostemon suaveolens* | Swamp box | 2 | Leaves | 2.61 | 50.27 | 1.63 | -28.2 | 19.25 |
| *Lophostemon suaveolens* | Swamp box | 3 | Leaves | 2.84 | 50.53 | 2.08 | -29.1 | 17.80 |
| *Melaleuca quinquenervia* | Paperbark | 1 | Leaves | 1.31 | 53.87 | 2.55 | -34.2 | 41.06 |
| *Melaleuca quinquenervia* | Paperbark | 2 | Leaves | 0.94 | 53.63 | 1.23 | -32.5 | 56.88 |
| *Melaleuca quinquenervia* | Paperbark | 3 | Leaves | 0.84 | 56.11 | 2.25 | -34.3 | 67.05 |
| *Casuarina glauca* | Swamp oak | 1 | Leaves | 2.04 | 47.14 | 1.38 | -30.0 | 23.16 |
| *Casuarina glauca* | Swamp oak | 2 | Leaves | 1.20 | 48.12 | -0.96 | -32.1 | 40.24 |
| *Casuarina glauca* | Swamp oak | 3 | Leaves | 1.46 | 46.62 | 2.25 | -30.9 | 31.89 |

**Table S6 (xlsx file). Sequencing metadata, community composition, and metabolic capabilities of the tree bark and soil metagenomes analysed.**

**Table S7 (xlsx file). Taxonomy, coverage, and metabolic capabilities of the metagenome-assembled genomes analysed.**

**Table S8. Biological nitrogen fixation (BNF) rates (in ng N g^-1^ day^-1^) measured for the 24 different trees and averaged for the eight different species (shaded).** Nitrogen fixation rates measured using the **^15^**N_2_ assimilation method on triplicate bark samples from three different trees of eight species across different types of forest. Data show mean ± SE, minimum and maximum BNF rates for each individual tree on triplicate bark samples. Shaded rows show average values at species level. Species are grouped by type of forest: mangrove (*Avicennia marina*), heathland (*Banksia integrifolia*, *Acacia leiocalyx*), coastal wetland (*Lophostemon suaveolens*, *Casuarina glauca*, *Melaleuca quinquenervia*) and upland forests (*Eucalyptus propinqua* and *Eucalyptus siderophloia*). Control samples are triplicate autoclaved bark samples incubated with **^15^**N_2_ gas.

| **Species** | **Type of Forest** | **Tree** | **Mean BNF** | **SE BNF** | **Min BNF** | **Max BNF** |
| --- | --- | --- | --- | --- | --- | --- |
| *A. marina* | Mangrove | 1 | 4.3 | 2.0 | 0.3 | 6.9 |
| *A. marina* | Mangrove | 2 | 4.2 | 2.2 | 0.1 | 7.4 |
| *A. marina* | Mangrove | 3 | 4.9 | 1.0 | 3.5 | 6.9 |
| *A. leiocalyx* | Heathland | 1 | 10.6 | 2.4 | 6.0 | 13.9 |
| *A. leiocalyx* | Heathland | 2 | 7.9 | 3.8 | 1.1 | 14.4 |
| *A. leiocalyx* | Heathland | 3 | 4.8 | 1.8 | 1.3 | 7.0 |
| *B. integrifolia* | Heathland | 1 | 3.6 | 1.2 | 1.9 | 5.9 |
| *B. integrifolia* | Heathland | 2 | 9.5 | 3.0 | 5.72 | 15.5. |
| *B. integrifolia* | Heathland | 3 | 3.1 | 0.8 | 1.6 | 4.3 |
| *C. glauca* | Coastal wetland | 1 | 11.0 | 1.4 | 8.2 | 13.0 |
| *C. glauca* | Coastal wetland | 2 | 21.9 | 12.4 | 7.2 | 46.5 |
| *C. glauca* | Coastal wetland | 3 | 20.0 | 8.6 | 5.6 | 35.2 |
| *L. suaveolens* | Coastal wetland | 1 | 8.2 | 3.3 | 2.5 | 14.0 |
| *L. suaveolens* | Coastal wetland | 2 | 17.0 | 8.3 | 0.7 | 27.5 |
| *L. suaveolens* | Coastal wetland | 3 | 13.6 | 2.0 | 10.6 | 17.5 |
| *M. quinquenervia* | Coastal wetland | 1 | 14.4 | 2.1 | 10.6 | 17.9 |
| *M. quinquenervia* | Coastal wetland | 2 | 38.2 | 27.5 | 9.3 | 93.1 |
| *M. quinquenervia* | Coastal wetland | 3 | 11.3 | 4.2 | 6.9 | 19.7 |
| *E. propinqua* | Upland | 1 | 6.5 | 0.2 | 6.1 | 6.9 |
| *E. propinqua* | Upland | 2 | 2.0 | 0.3 | 1.5 | 2.4 |
| *E. propinqua* | Upland | 3 | 2.9 | 0.6 | 2.1 | 4.1 |
| *E. siderophloia* | Upland | 1 | 1.3 | 0.9 | 0.1 | 3.0 |
| *E. siderophloia* | Upland | 2 | 4.8 | 1.0 | 3.3 | 6.6 |
| *E. siderophloia* | Upland | 3 | 2.8 | 1.4 | 0.3 | 5.3 |
| *A. marina* | Mangrove | Average | 4.4 | 0.9 | 0.1 | 7.4 |
| *A. leiocalyx* | Heathland | Average | 7.8 | 1.6 | 1.1 | 14.4 |
| *B. integrifolia* | Heathland | Average | 5.4 | 1.4 | 1.6 | 15.5 |
| *E. propinqua* | Upland | Average | 3.8 | 0.7 | 1.5 | 6.9 |
| *E. siderophloia* | Upland | Average | 3.0 | 0.8 | 0.1 | 6.6 |
| *L. suaveolens* | Coastal wetland | Average | 12.9 | 2.9 | 0.7 | 27.5 |
| *C. glauca* | Coastal wetland | Average | 17.6 | 4.7 | 5.6 | 46.5 |
| *M. quinquenervia* | Coastal wetland | Average | 21.3 | 9.1 | 6.9 | 93.1 |
| Control | Control | Average | 1.0 | 0.3 | 0.6 | 1.4 |

**Table S9. Biological nitrogen fixation (BNF) rates in *M. quinquenervia* bark under methane addition and methanotroph inhibition.** BNF rates (ng N g⁻¹ day⁻¹) were measured using the ^15^N_2_ assimilation method on triplicate bark samples over one- and two-day incubations. Treatments included: (1) stem gas collected from the same tree using a gas chamber, (2) stem gas with methane (0.26% CH_4_), and (3) stem gas with difluoromethane (2% DFM), a methanotroph inhibitor. δ^15^N values represent ^15^N enrichment in bark. Controls are triplicate autoclaved bark samples. Sample size n = 3.

| **Treatment** | **Incubation time (days)** | **Replicate** | **δ^15^N (‰)** | **BNF rate (ng N g^-1^ day^-1^)** |
| --- | --- | --- | --- | --- |
| Stem gas + CH_4_ | 1 | 1 | 2.94 | 144.63 |
| Stem gas + CH_4_ | 1 | 2 | 3.52 | 133.98 |
| Stem gas + CH_4_ | 1 | 3 | 5.31 | 202.56 |
| Stem gas | 1 | 1 | 1.86 | 63.40 |
| Stem gas | 1 | 2 | 1.58 | 70.23 |
| Stem gas | 1 | 3 | 1.92 | 70.53 |
| Stem gas +DFM | 1 | 1 | 0.96 | 23.61 |
| Stem gas +DFM | 1 | 2 | 0.89 | 32.32 |
| Stem gas +DFM | 1 | 3 | 1.72 | 99.17 |
| Stem gas + CH_4_ | 2 | 1 | 7.06 | 169.86 |
| Stem gas + CH_4_ | 2 | 2 | 5.76 | 189.43 |
| Stem gas + CH_4_ | 2 | 3 | 9.00 | 259.51 |
| Stem gas | 2 | 1 | 1.85 | 31.61 |
| Stem gas | 2 | 2 | 1.90 | 32.72 |
| Stem gas | 2 | 3 | 2.25 | 60.67 |
| Stem gas +DFM | 2 | 1 | 0.49 | 2.02 |
| Stem gas +DFM | 2 | 2 | 0.48 | 1.65 |
| Stem gas +DFM | 2 | 3 | 0.50 | 1.92 |
| Control | 2 | 1 | 0.62 | 6.46 |
| Control | 2 | 2 | 1.05 | 25.03 |
| Control | 2 | 3 | 1.02 | 17.56 |

**Table S10. Macronutrients and micronutrients of the bark and soil samples from all the species sampled based on the 2024 campaign.** Elemental analysis of bark and soil samples from eight Australian tree species sampled on the 2024 campaign and paired with the ^15^N_2_ labelling assays and analysed using LECO. Total carbon (C%), total nitrogen (%N), carbon-to-nitrogen ratios (C/N) and concentrations of molybdenum (Mo), vanadium (V), phosphorus (P), nickel (Ni), iron (Fe) and copper (Cu) in mg kg^-1^ are shown. Mo and V are important cofactor for molybdenum- and vanadium- containing nitrogenases, respectively.

| **Tree species** | **Common name** | **Sample type** | **Total C (%)** | **Total N (%)** | **C/N** | **Mo (mg/kg)** | **V (mg/kg)** | **P (mg/kg)** | **Ni (mg/kg)** | **Fe (mg/kg)** | **Cu (mg/kg)** |
| --- | --- | --- | --- | --- | --- | --- | --- | --- | --- | --- | --- |
| *Eucalyptus siderophloia* | Iron bark | bark | 53.00 | 0.25 | 212 | 0.11 | 0.12 | 40.00 | 1.48 | 126.00 | 3.05 |
|  |  | soil | 3.67 | 0.18 | 20.4 | 0.24 | 19.7 | 134 | 1.3 | 8912 | 2.47 |
| *Casuasina glauca* | Swamp oak | bark | 46.70 | 0.60 | 77.80 | 0.15 | <0.1 | 95.00 | 0.18 | 164 | 1.67 |
|  |  | soil | 2.86 | 0.15 | 19.1 | <0.1 | 1.2 | 63 | 0.5 | 421 | 0.86 |
| *Melaleuca quinquenervia* | Paperbark | bark | 58.50 | 0.28 | 209 | <0.1 | 0.1 | 61 | 1.76 | 158 | 0.86 |
|  |  | soil | 28.40 | 0.98 | 29 | 0.53 | 4.2 | 371 | 2.5 | 1736 | 6.64 |
| *Lophostemon suaveolens* | Swamp box | bark | 49.90 | 0.46 | 108 | <0.1 | 0.31 | 114 | 0.75 | 315 | 3.21 |
|  |  | soil | 5.66 | 0.29 | 19.5 | 0.33 | 5.1 | 139 | 2 | 1449 | 1.61 |
| *Banksia integrifolia* | Coastal Banksia | bark | 51.10 | 0.24 | 213 | 0.1 | <0.1 | 65 | 0.22 | 129 | 2.47 |
|  |  | soil | 7.37 | 0.26 | 28.3 | <0.1 | 1 | 66 | 0.4 | 447 | 1.72 |
| *Avicennia marina* | Grey Mangrove | bark | 48.00 | 0.33 | 145 | <0.1 | <0.1 | 564 | 0.18 | 80 | 1.27 |
|  |  | soil | 2.22 | 0.16 | 13.9 | 2.7 | 24.4 | 321 | 4.6 | 18931 | 6.23 |
| *Eucalyptus propinqua* | Small Fruited Grey Gum | bark | 43.30 | 0.21 | 206 | <0.1 | <0.1 | 126 | 0.25 | 37 | 1.92 |
|  |  | soil | 5.70 | 0.29 | 19.7 | 0.36 | 26.9 | 226 | 1.3 | 15207 | 3.75 |
| *Acacia leiocalyx* | Black wattle | bark | 48.30 | 0.74 | 65.3 | <0.1 | 0.13 | 322 | 0.24 | 114 | 1.87 |
|  |  | soil | 3.17 | 0.18 | 17.6 | 0.16 | 6.3 | 59 | 0.9 | 2926 | 2.4 |

**Table S11. Comparison of statistical models used for analysis of biological nitrogen fixation rates.** Five models were compared to determine the most appropriate statistical approach for analysing N fixation rates across species. Models included: Linear mixed-effects model (LME) with Box-Cox transformation (λ = 0.222) and tree as random effect, ANOVA with the same Box-Cox transformation and tree as fixed effect, Generalised linear model (GLM) with Gamma distribution and log link without random effects, Generalised linear mixed model (GLMM) with log link Gamma distribution and tree as random effect, GLM with tree nested into species and log link Gamma distribution. Model selection based on Akaike Information Criterion (AIC) and Bayesian Information Criterion (BIC), with ΔAIC representing the difference from the best fitting model (LME). The LME was the best fitting model, achieving normality of residuals with the lowest AIC. Despite tree being included as a random effect, the variance was negligible (SD = 5.7 x 10^-5^). Generalised linear models showed substantially worse fit (ΔAIC > 180).

| **Model** | **AIC** | **BIC** | **LogLik** | **df** | **ΔAIC** | **Notes** |
| --- | --- | --- | --- | --- | --- | --- |
| LME  (λ = 0.22 transformed) | 268.35 | 292.43 | -123.17 | 11 | 0 | Best model, data normal after Box-Cox transformation. Random effect (tree) is negligible |
| ANOVA  (λ = 0.22 transformed) | 282.78 | 343.04 | -115.99 | 26 | 14.43 | Improved normality using the same transformation, tree as fixed effect |
| GLM (no random effects, log link) | 451.75 | 474.92 | -215.87 | 10 | 183.4 | Simplified model without random effect |
| GLMM (Gamma, log link) | 455.02 | 480.52 | -216.51 | 11 | 186.67 | Accounts for tree variance as random effect |
| GLM (nested fixed effects, log link) | 461.16 | 521.41 | -205.58 | 26 | 192.81 | Tree nested in species, ignores random variability |
